# DNA sequence-dependent positioning of the linker histone in a nucleosome: a single-pair FRET study

**DOI:** 10.1101/2020.12.07.414334

**Authors:** Madhura De, Mehmet Ali Öztürk, Sebastian Isbaner, Katalin Tóth, Rebecca C. Wade

**Affiliations:** Department of Biophysics of Macromolecules, German Cancer Research Center (DKFZ), 69120 Heidelberg, Germany; Molecular and Cellular Modelling Group, Heidelberg Institute for Theoretical Studies (HITS), 69118 Heidelberg, Germany; Faculty of Biosciences, Heidelberg University, 69120 Heidelberg, Germany; Centre for Biological Signalling Studies (BIOSS) and Centre for Integrative Biological Signalling Studies (CIBSS), University of Freiburg, 79104 Freiburg, Germany; Center for Molecular Biology (ZMBH), DKFZ-ZMBH Alliance, Heidelberg University, 69120 Heidelberg, Germany; Interdisciplinary Center for Scientific Computing (IWR), 69120 Heidelberg, Germany; Research Institute of Molecular Pathology (IMP), Vienna BioCenter (VBC), Campus-Vienna-Biocenter 1, 1030 Vienna, Austria

**Keywords:** nucleosome, linker histone, single-pair FRET, linker DNA sequence, A-tract

## Abstract

Linker histones (LH) bind to nucleosomes with their globular domain (gH) positioned in either an on- or an off-dyad binding mode. Here, we study the effect of the linker DNA (L-DNA) sequence on the binding of a full-length LH, *Xenopus laevis* H1.0b, to a Widom 601 nucleosome core particle (NCP) flanked by two 40 bp long L-DNA arms, by single-pair FRET spectroscopy. We varied the sequence of the 11 bp of L-DNA adjoining the NCP on either side, making the sequence either A-tract, purely GC, or mixed, with 64% AT. The labelled gH consistently exhibited higher FRET efficiency with the labelled L-DNA containing the A-tract, than that with the pure-GC stretch, even when the stretches were swapped. However, it did not exhibit higher FRET efficiency with the L-DNA containing 64% AT-rich mixed DNA when compared to the pure-GC stretch. We explain our observations with a model that shows that the gH binds on-dyad and that two arginines mediate recognition of the A-tract via its characteristically narrow minor groove. To investigate whether this on-dyad minor groove-based recognition was distinct from previously identified off-dyad major groove-based recognition, a nucleosome was designed with A-tracts on both the L-DNA arms. One A-tract was complementary to thymine and the other to deoxyuridine. The major groove of the thymine-tract was lined with methyl groups that were absent from the major groove of the deoxyuridine tract. The gH exhibited similar FRET for both these A-tracts, suggesting that it does not interact with the thymine methyl groups exposed on the major groove. Our observations thus complement previous studies that suggest that different LH isoforms may employ different ways of recognizingff AT-rich DNA and A-tracts. This adaptability may enable the LH to universally compact scaffold-associated regions and constitutive heterochromatin, which are rich in such sequences.

**Statement of Significance:** Linker histones (LHs) associate with the smallest repeat unit of chromatin, the nucleosome. They have been observed to have affinity for AT-rich DNA, which is found in constitutive heterochromatin and scaffold-associated regions (SAR), which could explain how the LHs can compact such parts of the chromatin. How the LH recognizes such sequences is poorly understood. Using single-pair FRET and modelling, we provide experimental evidence of DNA-sequence-induced changes in the orientation of a LH bound to a nucleosome, and thereby reveal a new mechanism by which the LH can recognize A-tract sequences that are abundantly present in the SAR. Our results show that, depending on how the LH associates with the nucleosome, it can employ more than one mechanism to recognize AT-rich DNA.

## Introduction

The genetic material of cells consists of nucleoprotein polymers which constitute chromatin, the smallest repeat unit of which is the nucleosome. The nucleosome consists of a nucleosome core particle (NCP) flanked by 20-60 bp long linker-DNA (L-DNA) arms on either side. The NCP is composed of a core histone octamer wrapped by 1.65 turns (or about 150 bp) of DNA. Linker histones (LH), lysine-rich proteins of about 200 residues (1), bind to nucleosomes. They have a tripartite structure: a short, flexible N-terminal domain (NTD), a highly conserved and structured globular domain (gH) of about 70 to 80 residues, and a long, disordered C-terminal domain (CTD or ‘tail’). Being highly positively charged, the LH, along with the core histones, is well-suited to neutralizing the negatively charged, mutually repulsive DNA strands and aiding in the packaging of the chromatin. Indeed, the major function of the LH is chromatin condensation (2) and it has been observed to play a crucial role in the formation of higher-order chromatin structures (3). LH-mediated compaction of chromatin results in the silencing of genes and repetitive sequences (4, 5).

The location of the LH on the nucleosome has been widely studied both experimentally and computationally (6, 7). X-ray crystal structures of nucleosome-LH complexes for *Gallus gallus* H5 (an H1 variant) (8, 9) and *Xenopus laevis* H1.0b (10) LHs display an ‘on-dyad’ binding mode in which the gH binds at the centre of the stretch of core or NCP DNA that is bounded by the entering and exiting L-DNA arms. A second possible binding mode, referred to as ‘off-dyad’, has been observed by NMR (10) and cryo-EM (11). In the off-dyad binding mode, the gH binds in the region bounded by the dyad axis (the axis passing through the nucleosome, equidistant from the L-DNA entry and exit points) and either the entering or the exiting L-DNA. Apart from these two observed locations, the gH has been observed to adopt different orientations on the nucleosome (8, 10–12). From Brownian dynamics simulations, Öztürk *et al*. (2018) (13) found that the gH can associate with nucleosomes in a range of configurations, with the configuration adopted depending on the sequences of the gH and the DNA as well as post-translational modifications. It is important to note that these binding modes of the LH refer specifically to the gH and that the CTD is highly flexible. Using FRET, Caterino and Hayes (2011) (14) found that the highly disordered CTD becomes locally structured in the presence of nucleosomes. Fang et al. (2016) (15) observed that the ordering of the CTD in the presence of mononucleosomes is distinct from that observed in the presence of nucleosomal arrays. Cryo-EM structural studies (11) showed that whereas the gH bound on-dyad, the CTD associated with one of the L-DNA arms. The local ordering of the CTD in the presence of nucleosomes could be a result of electrostatic interactions between the negatively charged L-DNA and the positively charged CTD, as observed by Luque *et al*. (2014) (16) from mesoscale modeling at different salt concentrations. On the other hand, using nanosecond FCS (fluorescence correlation spectroscopy) measurements, Heidarsson et al. (2020) (17) observed that the CTD remains highly disordered upon binding mononucleosomes.

In-depth experimental and computational studies have pinpointed the amino acid residues in the gH that contribute to DNA binding (6, 18–20) and to the two different binding modes, on- and off-dyad (21). However, the possible contributions of the linker DNA (L-DNA) sequence to the LH binding mode and orientation have largely remained unclear.

Previous experimental and computational studies (9, 10, 18, 20, 22, 23), and 3D structures (PDB ids: 5NL0 (12), 4QLC (8), 5WCU (9), 7K5X (23)) of nucleosomes with the gH bound in an on-dyad position, have shown that the gH has three DNA contacting surfaces (shown in Fig. 1B): positively charged residues on the α3 helix (labelled Zone A), and the loop 1 and β2 (Zone B) interact with the L-DNA whereas hydrophobic residues on the beta-hairpin loop (Zone C) interact with the dyad DNA.

**Figure 1:**
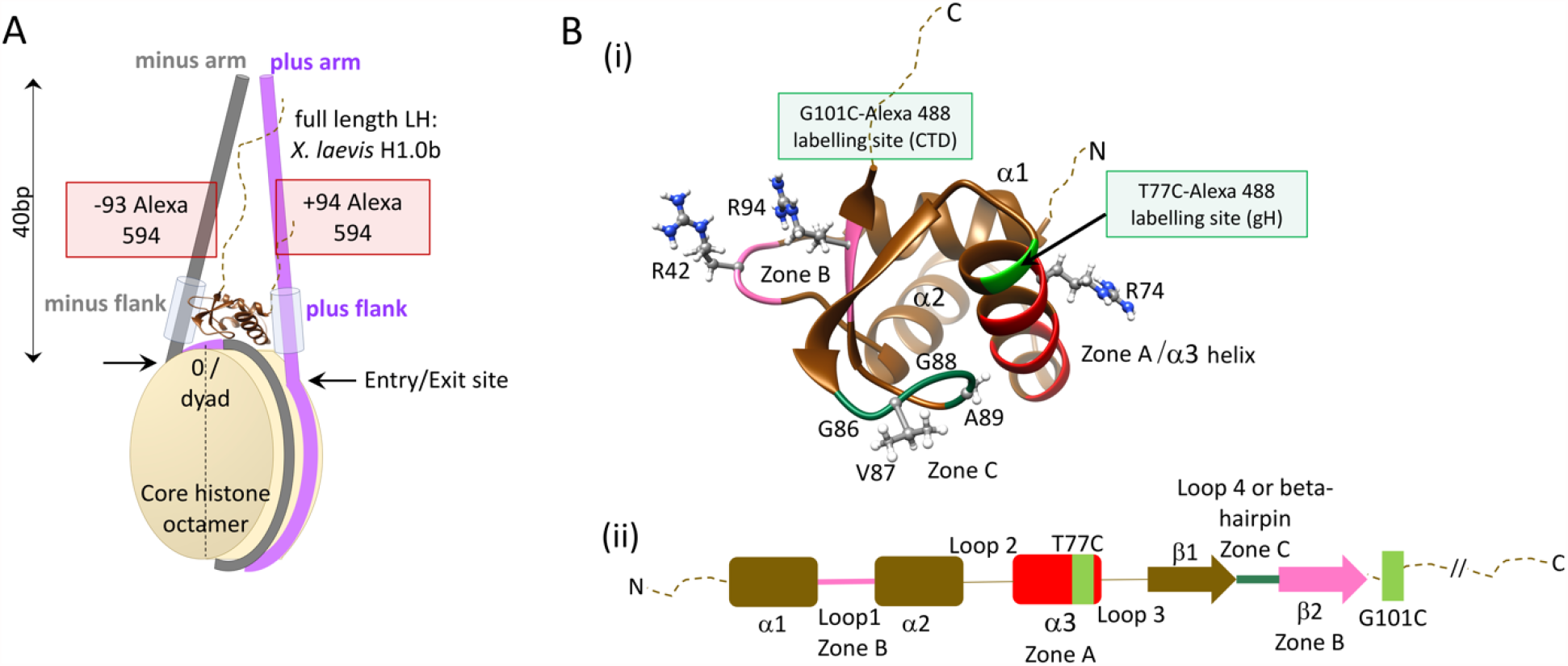
Architecture of the nucleosomes and linker histone studied. **A**. Schematic diagram of the nucleosome reconstituted from 226 bp non-palindromic Widom 601 DNA. The dyad base pair is numbered 0. The minus arm of the L-DNA is coloured grey and the plus arm is coloured purple. The Alexa 594 labels are attached on thymines -93 and +94, each 20 bp from the ends of the L-DNA arms. **B (i)**. Structure of the linker histone with the gH shown in cartoon representation with the positions of the two Alexa 488 labels at T77C and G101C on the CTD. The three DNA-proximal zones when the LH binds on-dyad are shown: A (red) or the α3 helix and B (pink) comprising of loop L1 and the beta sheet β2, interact with the two L-DNA arms. The beta-hairpin or zone C (deep green) interacts with the core or NCP DNA. Important residues are shown in ball-and-stick representation: R74 (Zone A), R42 and R94 (Zone B), and G86, V87, G88 and A89 (Zone C hydrophobic patch). **(ii)**. Schematic representation of the LH sequence and secondary structure with the DNA-proximal zones indicated. Dotted lines represent the disordered N- and C-terminal domains. The two attachment sites of the Alexa 488 donor dye are indicated in green.

Although the LH has not been reported to have any consensus DNA binding sequence, preferences for certain sequences have been noted. A number of in-vitro studies on the binding of LH to naked DNA (24, 25) showed an affinity of the LH for AT-rich sequences. Evidence of the LH having an affinity for AT- and A-tract-rich <u>s</u>caffold <u>a</u>ttachment <u>r</u>egions (SAR) (26) and for satellite sequences (27) led to the proposition that the LH has an affinity for not just AT-rich sequences but specifically for A-tract regions (sequences with a stretch of 4 to 6 or more adenine bases (28)). This was further confirmed by Roque et.al. (2004) (29) who showed that LH selectively bound to A-tract-rich SARs in the presence of competing DNA sequences having an AT content of 75% but totally lacking in A-tracts. Recent tunneling electron microscopic images and sedimentation studies on A-tract-containing nucleosomal arrays also showed strong compaction in the presence of LHs (30). The affinity of LHs for AT-rich sequences has been attributed to both the gH (31, 32) and also to the CTD alone (29).

The biological relevance of the preference of LHs for AT-rich or A-tract sequences was pointed out in a review by Zlatanova *et al*. (1991) (24). They stated that a preference of LHs for AT-rich sequences (including A-tracts) might aid the LH-mediated compaction of AT-rich heterochromatin. On the other hand, while LHs have been shown to have affinity for A-tract sequences, the opposite has been observed for GC-rich sequences (23, 27, 33).

The aim of the current study is to observe the effects of the L-DNA sequence on the positioning and orientation of the LH in the nucleosome, and to pinpoint the underlying reasons for the affinity of the LH for specific DNA sequences, such as A-tracts. We used single-pair FRET to probe the location and orientation of the full-length *Xenopus laevis* LH, H1.0b, on mononucleosomes reconstituted from recombinant, full-length, *X. laevis* core histones and non-palindromic Widom 601 sequences flanked by 40 bp L-DNA arms. We modified a stretch of 11 bp on each L-DNA, adjacent to the points where the DNA enters and exits the nucleosome (the ‘entry-exit’ sites). Since these stretches flank the nucleosome core, we refer to them as ‘flanks’. We study the effects of a pure GC stretch, a mixed sequence (64% AT-rich), and A-tracts pairing with thymine as well as deoxy-uridine, on the position and orientation of the LH. Based on FRET-derived distances, we suggest how A-tracts are recognized by the LH. Our model shows that the gH is oriented towards A-tracts so as to position R42 and R94 close to the narrow minor groove of A-tract DNA. Recognition of A-tracts via their characteristic narrow minor grooves by proteins containing positively charged amino acid residues, such as arginines, has been well documented (28, 34, 35). We suggest that this very same mechanism could be responsible for the specific recognition of A-tracts by the LH.

## Materials and Methods

### Preparation of labelled DNA

226 bp unlabelled and singly labelled DNA was used for reconstituting mononucleosomes. Labelled DNA was produced using PCR, from either the pGEM3z vector for the MG construct (see Supporting Material: ‘Sequences and Labelling Positions’), or from 226 bp templates (Biolegio, Nijmegen, The Netherlands). All the DNA sequences studied contain the strongly positioning sequence, Widom 601 (36). Primers used to amplify the DNA (IBA Lifesciences, Goettingen, Germany) contained the fluorophore Alexa 594, covalently attached via an amino-dT C6 linker. For the TU construct, the stretch of 11 deoxyuridines was incorporated in the reverse primer (Biolegio, Nijmegen, The Netherlands), (Supporting Material: ‘Sequences and Labelling Positions’). PCR was performed using Taq-polymerase containing master mix (Thermo Scientific). The final 226 bp product consisted of the highly positioning, non-palindromic Widom 601 sequence at the centre, bounded by two equally long (∼40 bp) L-DNA. The fluorophores were attached 20 bp from the ends of the DNA, on one or the other L-DNA arm (Fig. 1a). The DNAs were purified from PCR products using Gen-Pak FAX HPLC (Waters) and gel electrophoresis was performed to check for the correct size of the product.

The central base pair of the DNA was numbered 0. Bases to the left were assigned negative numbers and bases to the right were assigned positive numbers. The labelling positions on the L-DNA arms were either +94 or -93. Based on the flank sequences (i.e. the stretches of 11 bp DNA flanking the central non-palindromic Widom 601), the 226 bp DNAs were named AG (A-tract minus flank, purely GC plus flank), GA (purely GC minus flank, A-tract plus flank), MG (mixed sequence minus flank, purely GC plus flank), GM (purely GC minus flank, mixed sequence plus flank), and TU (A(T)-tract minus flank, A(U)-tract plus flank) (Supporting Material: ‘Sequences and Labelling Positions’, and Table 1). Nucleosomes reconstituted from these DNA sequences were named accordingly.

**Table 1:**
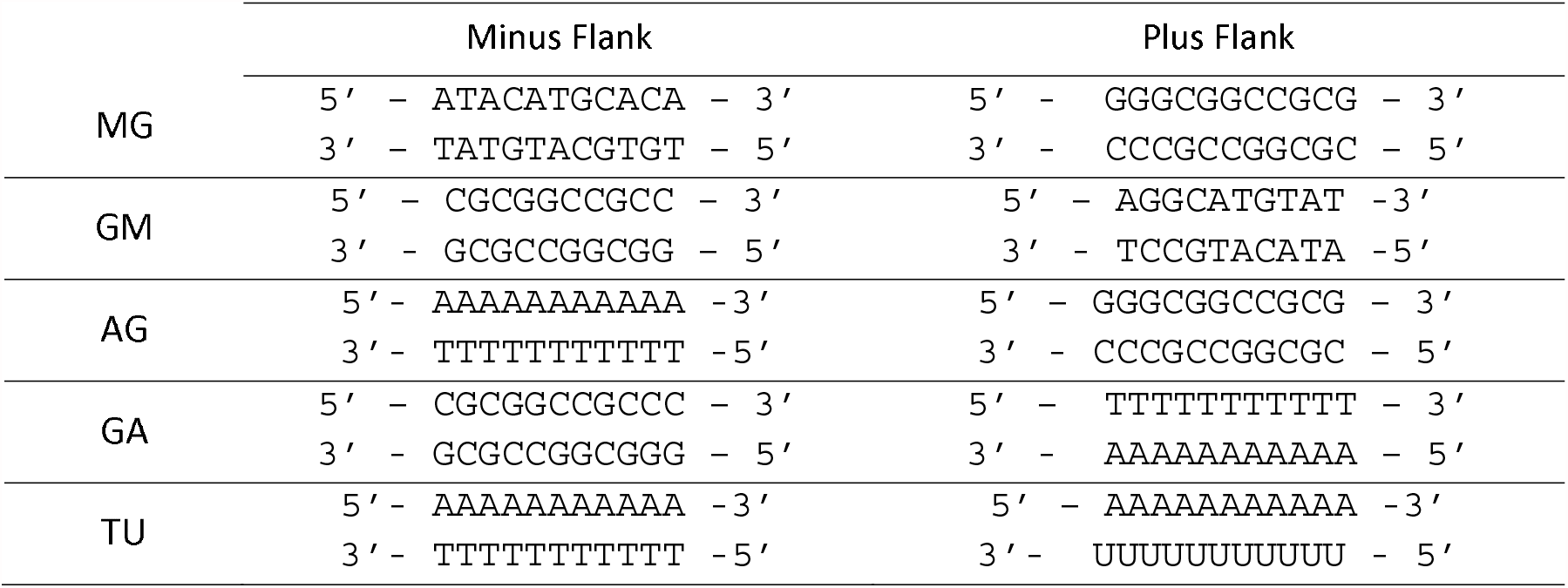
The ‘flank’ sequences of the different nucleosome constructs studied. ‘U’ denotes deoxyuridine (dU) in the plus L-DNA of the TU construct. See Supporting Materials ‘Sequences and Labelling Positions’ for the full DNA sequences.

### Preparation and purification of LH

Full-length, recombinant LH from *Xenopus laevis*, H1.0b, was used. Unlabeled, wild-type LH was used with either unlabeled nucleosomes or nucleosomes labelled only on the DNA (acceptor-only nucleosomes). We labelled the LH with Alexa 488 (Thermo Fischer Scientific), the donor fluorophore, either on the CTD or on the gH. For labelling, cysteine residues were first incorporated by site-directed mutagenesis in the cysteine-free, wild-type LH. The plasmid for the gH label (with the T77C mutation), and the protein for the CTD label (with the G101C mutation) were kindly provided by Prof. Jeffrey Hayes (University of Rochester Medical Center). the plasmid containing the linker histone T77C mutant gene was transformed into and amplified in *E*.*coli* XL-1 Blue competent cells. Amplified plasmids were purified and retransformed to *E*.*coli* BL21(DE3) cells for protein expression, with IPTG induction as described in reference (37). The bacterial pellet was sonicated (Branson Sonifier 250) in a lysis buffer (50 mM Tris-HCl, pH 7.5, 100 mM KCl, 1 mM EDTA, 1 mM phenylmethanesulphonylfluoride, 0.1 % volume/volume Nonidet P-40). After multiple rounds of washing the bacterial pellet, the supernatant was concentrated and simultaneously buffer exchanged to an unfolding buffer (7 M guanidium hydrochloride, 20 mM Tris.HCl, pH 7.5), using Vivaspin 2 columns (10 kDa molecular weight cutoff, Sartorius). The resulting, concentrated protein solution was passed through a size exclusion chromatography column, Sephacryl S-200 (pre-equilibrated by SAU-1000 buffer: 7 M deionized urea, 20 mM sodium acetate, pH 5.2, 1 mM EDTA, 1 M KCl), at a flow rate of 3 ml / min. The eluted protein fractions were concentrated and simultaneously buffer-exchanged to SAU-50 buffer (7 M deionized urea, 20 mM sodium acetate, pH 5.2, 1 mM EDTA, 50 mM KCl) using Vivaspin 20 columns (10 kDa molecular weight cutoff, Sartorius). This step was followed by cation exchange chromatography using a Mono-S-HR 10/10 FPLC column pre-equilibrated with SAU-50 and SAU-1000 buffers. The protein was eluted with SAU-1000 buffer. Protein eluate was concentrated and refolded by overnight dialysis at 4°C into 2 M NaCl, 10 mM Tris-HCl, 0.1 mM EDTA, pH 7.5.

Wild type, recombinant, full-length X. laevis core histones were purchased from Planet Protein (Colorado State University, USA), assembled into octamers, and purified as described previously (38, 39).

### LH labelling

The LH was labelled with Alexa 488 C5 maleimide (Thermo Fischer Scientific) on the cysteine residue either on the gH (position T77C) or the CTD (position G101C) (see Figures S1 and S2 in Supporting Materials). Unlabelled, cysteine-containing protein was unfolded in a denaturing buffer containing 7 M guanidine hydrochloride, 10 mM dithiothreitol, 20 mM Tris-HCl, at pH 7.5. Tris (2-carboxyethyl) phosphine was added as a reducing agent, at a concentration ten times higher than the protein concentration, to prevent the formation of protein dimers through a disulfide bridge. Alexa 488 C5 maleimide dye (Thermo Fischer Scientific) was dissolved in N’N’-dimethyl formamide, and added in three steps every hour, to the protein shaking in the dark at room temperature, to achieve a final dye concentration of ten times the number of moles of protein. This mix was stored at 4°C overnight. The maleimide reaction was stopped on the following day, by adding L-cysteine at the same concentration as the dye molecules. The labelled protein was dialysed into an unfolding buffer (urea 7 M, sodium acetate 20 mM, pH 5.2, EDTA 1 mM, KCl 1 M). To remove excess, unbound dyes, the protein was passed through a sephacryl S200 column. Finally, the protein was refolded by multiple dialysis steps in 2 M NaCl TE (10 mM <u>T</u>ris-HCl, 0.1 mM <u>E</u>DTA) buffer (pH 7.5). The Alexa 488 dye at the labelling position on the gH (T77C) was modeled in the crystal structure 5NL0 (12) using the software FRET Positioning and Screening (40) to ascertain that it did not clash with either the L-DNA or the dyad-DNA on the NCP. The dye position G101C on the CTD has been used in previous studies (12, 15, 41, 42). Since this position is only 6 residues away from the gH domain and not further downstream in the intrinsically disordered CTD, we expected to obtain a good signal to noise ratio from this position.

### Nucleosome reconstitution

Nucleosomes were reconstituted from recombinant *Xenopus laevis* core histones, 226 bp labelled or unlabelled DNA, and full length, recombinant, *Xenopus laevis* LH that were either labelled or unlabelled. Reconstitution involved reducing the salt concentration by a two-step dialysis method, using Slide-A-Lyzer MINI Dialysis unit (Thermo Fischer) having a cut-off of 7 kDa. DNA and core histone octamers were mixed in a molar ratio of 1 : 1.65 (39) in TE buffer at with 2 M NaCl. The first step of the dialysis, performed for four hours at 4°C, was against 0.6 M NaCl in TE. This lowered the NaCl concentration of the solution containing DNA and core histones from 2 M to 0.6 M, enabling reconstitution of mononucleosomes. The LH was added at this stage (43), at a molar ratio of 1.6 : 1 mole DNA (44), taking care to have a dye stoichiometric ratio (measured in ALEX, see section Single-pair FRET (Materials and Methods) and Supporting Methods: ‘Obtaining inter-dye distances from single-pair FRET and bulk spectroscopy’) of 1 : 1 (S = 0.5) donor (on LH) and acceptor (on DNA), while avoiding aggregation due to excess LH. Dialysis was performed against TE buffer without NaCl at 4°C overnight, until the NaCl concentration of the DNA and protein mix was lowered to 25 mM. All single-molecule and bulk spectroscopic measurements were performed at 25 mM NaCl in TE buffer. Nucleosome formation was checked by 6 % polyacrylamide gel electrophoresis (acrylamide : bisacrylamide 60 : 1) (Figure S3 in the Supporting Materials). Single labeled nucleosomes such as donor-only (Alexa 488 label on the LH) and acceptor-only (Alexa 594 label on the DNA, the LH remains unlabeled) samples were reconstituted in the same way as double labelled samples.

### Single-pair FRET

Single-pair FRET with <u>A</u>lternating <u>L</u>aser <u>Ex</u>citation (ALEX) (45, 46) experiments were performed on freely-diffusing, doubly labelled samples, using our in-house instrument described in (47) that was further modified (48) by the inclusion of the acceptor laser (561 nm, Cobolt Jive, Hübner Photonics) and an acousto-optic tunable filter (AOTFnC-VIS-TN, AA Optoelectronics) to switch between the two wavelengths (491 nm, Cobolt Calypso, Hübner Photonics, and 561 nm) on a microsecond timescale. The laser intensity at the objective, for measuring at single-molecule levels, was 40 μW. The light collected from the sample comprises a mix of fluorescence emitted from the donor and acceptor fluorophores. The emitted beam is split using a dichroic mirror (600DCXR, Omega Optical, Brattleboro, USA) to donor and acceptor signals which are sent through filters (donor channel: 520df40, acceptor channel: 610ALP, Omega Optical) to two avalanche photodiodes (SPAD-AQ-14, PerkinElmer Optoelectronics, Boston, Massachusetts, USA), see Figure S4 for a schematic of the setup adapted from (48). Although we obtained both the stoichiometry (proportion of the donor and acceptor dyes) and the proximity ratio by this procedure (see Figure S4, and in Lee *et al*. (2005) (46)), we extracted the proximity ratios of only the doubly labelled samples, i.e., having a stoichiometry range of 0.25 to 0.75. The proximity ratio is the ratio of the number of photons detected in the acceptor channel divided by the total number of photons detected in both the donor and acceptor channels upon excitation of the donor (46). The equation describing the proximity ratio is:

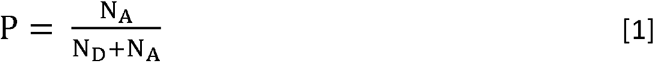

where N_A_ and N_D_ are the number of photons detected in the acceptor and donor channels respectively (see the section ‘Methods-Obtaining inter-dye distances from single-pair FRET and bulk spectroscopy’ in the Supporting Materials). Proximity ratio histograms were plotted using our in-house AlexEval software (48) with 50 bins, a bin width of 0.02, and ranging from -0.1 to 1.1. Multiple Gaussian peaks were fitted to the proximity ratio histograms. The proximity ratio is related to the FRET efficiency E, via an instrumentation detection factor γ (Proximity ratio = FRET efficiency when γ = 1), that we measured prior to sample measurements, using two FRET standards: 30 bp oligos labelled with Alexa 488 and Alexa 594 at a separation of 10 and 21 bp (Table S3 in the Supporting Material, γ factor calculation is described in Lee *et al*. (2005)(46)). Using bulk fluorescence and absorption measurements, we obtained a Förster distance, R_0_, of 52 Å, assuming a refractive index of 1.4 (see Table S1 for determination of R_0_ and Table S2 for a comparison with R_0_ values in the literature).

From the FRET efficiency, using R_0_, we obtained inter-fluorophore distances (Supporting Methods: ‘Obtaining inter-dye distances from single-pair FRET and bulk spectroscopy’ and Figure S5 in the Supporting Materials), which were used to model the nucleosomes AG, GA, and TU.

For the single-pair FRET measurements, we used 50 to 100 pM of doubly labelled samples, and added unlabelled samples to raise the total concentration of the samples to 300 pM. This procedure avoided spontaneous dissociation of the reconstituted nucleosomes (47). Measurements were done for 30 minutes in 25 mM NaCl 1xTE buffer, pH 7.5. The sample solution also contained 0.8 mM ascorbic acid for scavenging free radicals to prevent photobleaching of the dyes as described previously (38), and 0.01 mM nonidet-P40 (49) to stabilize the mononucleosomes.

### Modeling of the AG, GA, and TU nucleosomes

Nucleosome models were built to interpret the experimental data obtained from the AG, GA, and TU constructs (see Materials and Methods: Preparation of labelled DNA for the names of the constructs). The crystal structure from Bednar et al. (2017) (12) (PDB: 5NL0) and a recent cryo-EM structure from Zhou et al. (2021) (50) (PDB: 7K5X) were used as starting structures to build the models of the AG and the GA nucleosomes. The gH and additional proteins (scFv20 in the case of PDB: 7K5X) were removed from the original structures and the gH coordinates were saved separately. Using Chimera (51), the DNA sequences of both the starting structures were modified to that of the AG and GA constructs, using the ‘swapna’ command. The L-DNA arms in the models had a length of 23 bp and included the dye labelling positions 3 bp from the ends, and 20 bp from the NCP DNA. In contrast, the L-DNA arms of the nucleosomes in our experiments were 40 bp long and the dyes, positioned 20 bp from the NCP, were 20 bp from the ends. Although longer L-DNA of 40 bp was used in the experiments, the dye at the mid-point of the L-DNA did not give any distance information for the portions of the L-DNA beyond the dyes and extending to the L-DNA ends. We thus omitted these portions from our models by modelling nucleosomes with L-DNA of 23 bp length, including the labeling positions. The structures of the AG and GA nucleosomes, each with two L-DNA conformations (5NL0-type and 7K5X-type), were protonated using Chimera to optimize the hydrogen bond network. While keeping the core histone octamers fixed, the DNA was energy minimised in Chimera using the AMBER ff99bsc0 force field (52) to remove clashes or kinks in the DNA backbone that may have resulted during the sequence modification. The minimisation protocol was as follows: 100 steps of steepest descent energy minimisation (step size 0.02 Å) followed by 10 conjugate gradient steps (step size 0.02 Å). The 3DNA 2.0 web server (53) was used to check that no A-form DNA was present, and that the DNA was predominantly in the standard B-form geometry (over 90% of the DNA base steps were B-form after energy minimisation). To sample the L-DNA arm motions (detailed in the caption of Figure S6 in the Supporting Materials), the 5NL0-type structures of the AG and GA nucleosomes were subjected to elastic network normal mode analysis using the elNémo webserver (54) to generate a range of L-DNA arm openings. Two conformers showing the maximum L-DNA arm openings, mode 7 frame 51 and mode 8 frame 1, were selected. Thus, we obtained four different types of L-DNA arm-opening conformations for both the AG and the GA nucleosomes: the 5NL0-type, the 7K5X-type, the mode 7 frame 51, and the mode 8 frame 1 (Fig S6).

For the TU construct, the 5NL0 structure was used as the starting structure. The DNA was mutated in Chimera in the same way as for AG and GA, but the L-DNA dynamics were not sampled further. Hydrogens were added and the construct was energy minimised in the same way as AG and GA. DNA sequences such as A-tracts are known to impart an inherent bend in the DNA (28, 30, 55–57). To consider this possibility, the L-DNA arms of AG and GA were modeled separately using the cgDNA webserver, with Paramset+ 1 (58) (see Figure S6 in the Supporting Materials). New AG and GA nucleosomes were built by removing the existing L-DNA from the entry/exit sites bounding the core and placing the cgDNA modeled L-DNAs onto the entry/exit sites. We found that the extent of bending induced by the different flanking sequences was sampled by the four arm-opening conformations studied (Figure S6 C). The gH domains were separately mutated to the sequence of H1.0b with a cysteine at position 77, using the ‘swapaa’ command in Chimera. After mutation, the gH was minimised in Chimera, using the AMBER ff14SB force field (59), using the same minimisation protocol as used for minimising the nucleosome DNA. Then, using Chimera, the gH was repositioned onto the nucleosome structures in various orientations to generate an ensemble of configurations of the nucleosome-gH complexes. These ensembles were screened using the FRET Positioning and Screening software (40) (Fig S7, Table S4 in the Supporting Material), with experimentally-derived distances as the input and a R_0_ value of 52 Å.

## Results and Discussion

### The globular domain of the Xenopus laevis H1.0b shows a preference for A-tracts

In this study, we asked how the L-DNA sequence may affect the location and/or the orientation of the LH. Does the LH prefer an AT-rich L-DNA arm over a GC-rich arm? Does the LH specifically prefer A-tracts (one strand purely adenines) or simply any sequence having a higher AT content than GC content?

To address these questions, we constructed two types of nucleosomes: in the MG construct, the minus flank (11 bp stretch adjacent to the core DNA) preceding the Widom 601 core DNA has a mixed sequence with a purine-pyrimidine base-step and 64% AT content. The plus flank is purely GC (Table 1). In the AG construct, the mixed sequence of the minus flank was replaced with an A-tract. The plus flank was left intact. In the GM and the GA constructs, the minus and the plus flanks of MG and AG, respectively, were swapped.

The gH of the full-length LH was labelled with the donor (Alexa 488) dye at residue 77 (threonine mutated to cysteine prior to labelling), at the C-terminal end of helix α3 (see Fig. 1B). The DNA was labelled with the acceptor (Alexa 594) dye on either the minus arm (−93 T) or on the plus arm (+94 T) (Fig. 1A), generating two FRET pair combinations per construct.

In the MG construct, the gH dye was observed to show a higher proximity ratio for the +94 dye (Fig. 2A), than for the -93 dye, suggesting that the gH dye is closer to the purely GC plus flank than the 36% GC/64% AT minus flank. This observation is at odds with previous *in vivo* and *in vitro* studies (27, 33) that reported the LH to have lower affinity for GC-rich regions than AT-rich regions.

**Figure 2:**
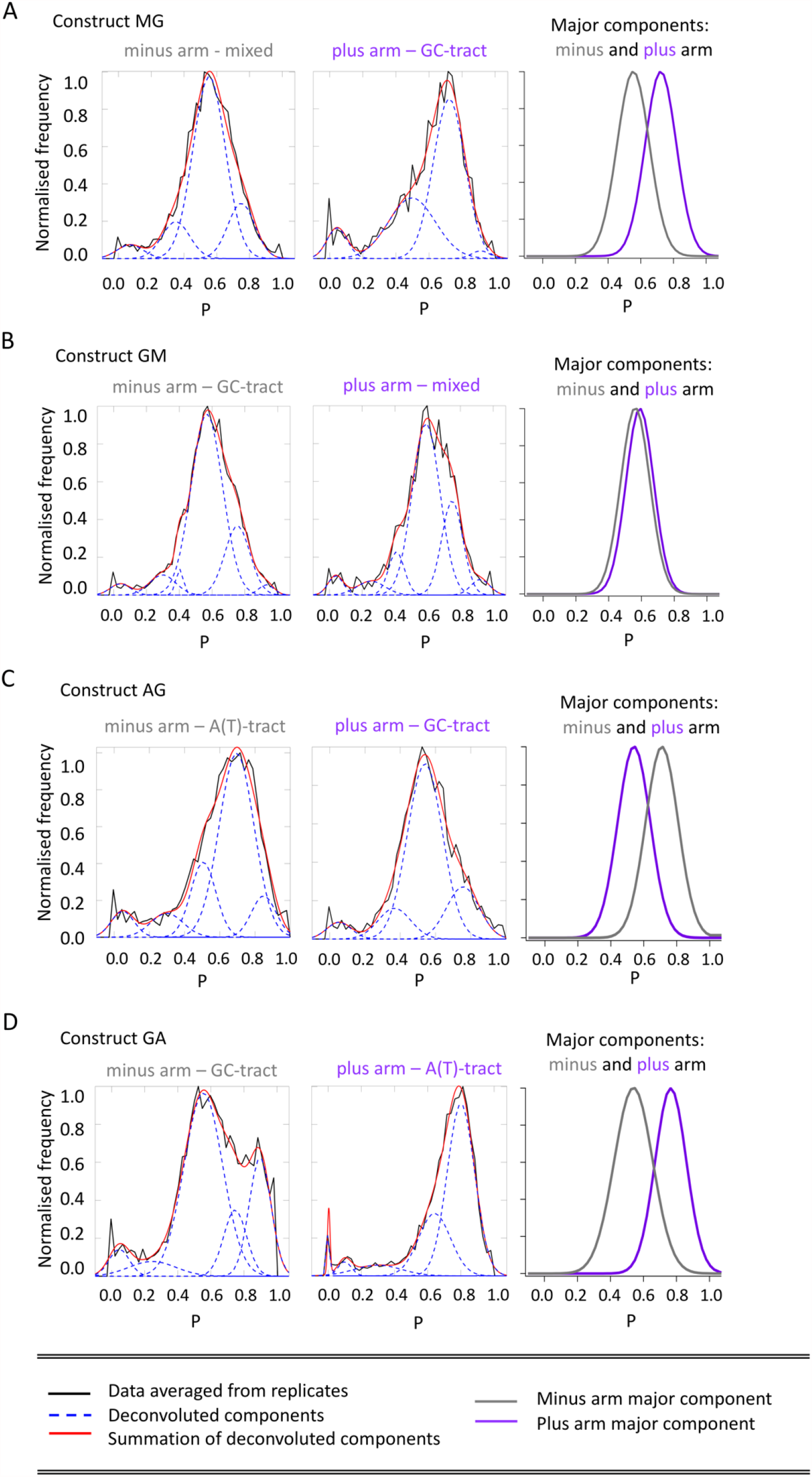
Proximity ratio (P) histograms for nucleosomes MG **(A)**, GM **(B)**, AG **(C)**, and GA **(D)** labelled on the plus (purple) or minus (grey) L-DNA arms and the gH site of the LH (T77C). From left to right for each of the nucleosomes A-D, the plots show fitted proximity ratio histograms of the minus arm, the plus arm, and the major population peaks from the minus arm (grey) and the plus arm (purple) (height normalised) shown on one plot for comparison. **(A)** The gH dye exhibits a higher proximity ratio for the GC-tract containing arm in the MG construct. **(B)** For the GM construct, the gH dye exhibit nearly equal proximity ratios for both the GC-tract containing arm and the mixed-DNA arm. **(C)** For the AG construct, the gH dye exhibits a higher proximity ratio for the A-tract containing arm and for the GA construct. **(D)** The major population peak in the minus arm histogram suggests that the gH exhibits a lower proximity ratio for the GC-tract than for the A-tract.

We thus built the GM construct as a control, with the minus and the plus flank of MG swapped. However, in the GM construct, the FRET profile of the gH dye is almost the same for the purely GC flank minus arm and the mixed flank plus arm (Fig. 2B, Table 2 and Figure S8, Table S5 in the Supporting Material). This observation suggests that the high FRET efficiency peak shown for the GC-flank in the MG construct could be due to the orientation of the gH on the MG nucleosome, and not due to a preference of the gH for GC-rich DNA.

**Table 2:**
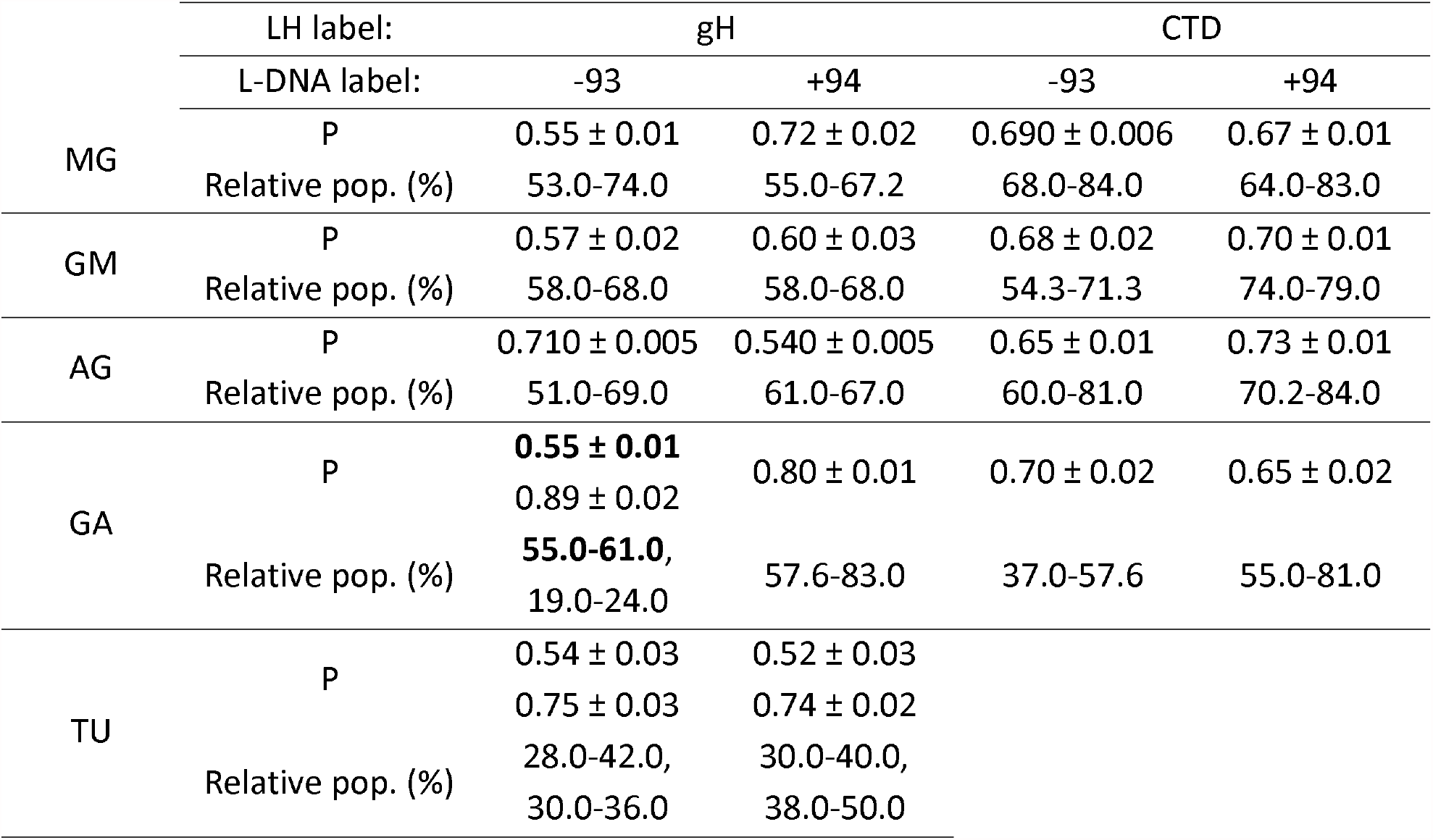
Proximity ratio (P) between the donor fluorophores on the LH (gH or CTD) and the acceptor fluorophores on the minus or the plus L-DNA arms of the five nucleosome constructs studied. If more than one population is consistently present, values for both are given, with those for the reproducibly major population in bold. The relative populations of the peaks are given as percentages. The ranges of the relative populations were obtained from three to six replicates (see Supporting Materials Tables S5 to S9).

We next considered whether the DNA sequence of the non-palindromic Widom 601, owing to its asymmetric, has any effect on the gH position or orientation. However, this possibility was ruled out, since the NCP DNA remained the same in the GM construct. The Widom 601 sequence was previously shown to unwrap asymmetrically by Ngo et al.(2015) (60), who also showed that the unwrapping was not caused by the DNA sequences at the entry/exit sites. Thus, had the asymmetric unwrapping of the Widom 601 nucleosome been a reason for the MG observation, swapping the flank sequences close to the entry/exit sites of the GM construct would have resulted in the same observation seen for MG. Secondly, if there was any effect of the GC-tract in bending the plus L-DNA closer to the gH, then we would have observed these effects for the minus arm in the GM construct as well. This too, was not the case. Thus, we suggest that the apparent preference of the gH for the plus arm of the MG construct is due to the orientation of the gH in this construct.

Despite using full-length core histone octamers, we cannot comment on the possible effects of the core octamer tails from our current data. Previous FRET studies (41, 61), computational studies (62, 63) and recent cryo EM structures (50) show that the core histone tails, specifically the H3 N-terminal and H2A C-terminal tails associate with the L-DNA at the flank regions. However, despite the symmetry of the core histone octamers, the tails associate with the flanks in an asymmetric fashion(61, 63), possibly due to the non-palindromic Widom 601 sequence. However, removal of core histone tails was found not to significantly affect the binding of an on-dyad positioned H5 gH domain (64). The occupancy of the two H3 tails from the MD simulation trajectories from the work of Shaytan et al. (2016) (63) suggests that the tails might not interfere with the binding site of an on-dyad positioned gH domain. However, since previous studies were performed on the standard non-palindromic Widom 601 sequence, it is unclear how the core histone tails interact with the L-DNA when the flank sequences are swapped, i.e., modifying the MG to the GM construct.

When the 64 % AT-rich mixed DNA flanks to which the gH showed no preference, were replaced by 11 bp A-tracts, the gH exhibited a higher proximity ratio to this flank than to the purely GC flank. The A-tract-containing minus arm of the AG construct (Fig 2C), and the plus arm in the GA construct (Fig 2D) consistently show a higher proximity ratio. Accordingly, the plus dye of the AG construct and the minus dye of the GA construct showed a lower proximity ratio (Fig 2C and 2D, Table 2, and Figure S9, Table S6 in the Supporting Material). We also observed a high proximity ratio peak (mean proximity ratio = 0.9, Table 2 and Table S6) for the minus dye in the GA construct. The relative area of this high FRET peak ranged from 19 to 24 % in the replicates whereas the dominant population (mean P = 0.55) comprised 55 to 61 % of the total area of the plot.

These observations suggest that: (a) the gH does not have a preference for pure GC-containing regions over a mixed DNA sequence, and (b) the gH does not have any preference for mixed DNA sequences, even if they have a higher AT content than GC. However, as shown by the AG and GA constructs, when the mixed 36 % GC / 64 % AT sequence of MG and GM is replaced by an A-tract, the gH consistently exhibits a higher proximity ratio with the A-tract flank. We next investigated the possible reasons behind the selective preference of the LH for A-tract regions.

### Arginines 42 and 94 in the globular domain of Xenopus laevis H1.0b mediate A-tract recognition

The inter-label distances of the major populations were derived from the proximity ratio histograms by a multi-step procedure based on both bulk and single-molecule spectroscopic measurements as described in Materials and Methods and Supporting Methods: ‘Obtaining inter-dye distances from single-pair FRET and bulk spectroscopy’. We derived a Förster distance of 52 Å for our samples and used it to calculate the FRET distances in the constructs AG and GA. Figure 3A and B panel (i) shows the dye to dye distance plots for the dominant populations obtained from the dyes on the plus (purple) and minus (grey) arms. For the distance histograms derived from the proximity ratio plots of each replicate, refer to Figure S13 and Table S10 in the Supporting Materials. In the AG construct, the gH dye is closer to the minus dye on the A-tract-minus arm with a mean distance of 44.2 ± 0.84 Å, and is about 50.0 ± 0.24 Å from the plus dye on the GC-tract plus arm (Fig 3A, Table 3). In the GA construct, the gH dye is 41.0 ± 0.74 Å from the plus dye on the A-tract plus arm and 49.0 ± 0.74 Å from the minus dye on the GC-tract minus arm (Fig 3B).

**Figure 3:**
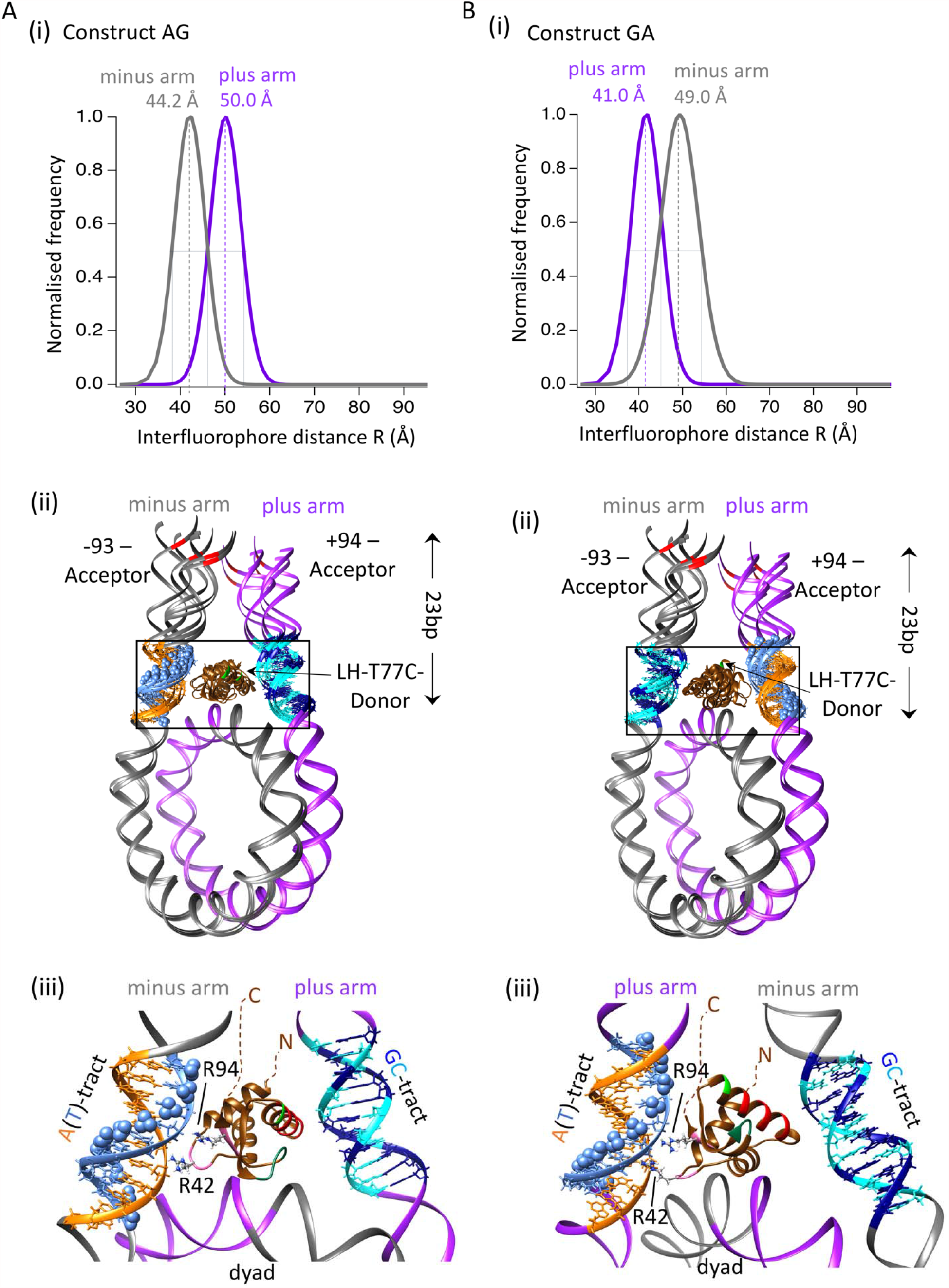
Models of the LH-nucleosome complexes consistent with the FRET data. Distance histograms of AG **(A)** and GA **(B)** nucleosomes **(i)** with their respective model ensembles **(ii)**, including close-up views **(iii)** showing the possible interactions between zone B (pink backbone) arginines and the A-tract minor grooves. Only the major population peaks are denoted in the distance plots (**Ai** and **Bi**). The mean distances of the peaks are given on the top of the peaks. The ‘representative’ models are those showing distances closest to the mean distance values (Table 3) obtained from experiments and are shown in detail in **(iii)**. The dotted lines schematically represent the disordered NTD and CTD. The flank sequences are shown in: orange (dA), light blue (dT), cyan (dC), and deep blue (dG). Thymine methyl groups are denoted by light blue spheres. **Biii** the view of the gH on the GA nucleosome is rotated 180 degrees from **Bii** so as to keep the A-tract on the plus arm on the left and the GC-tract on the minus arm on the right.

**Table 3:** Distance (R) or Proximity ratio (P) (in case of TU) between the dyes on the plus or minus arm and the gH (T77C site) from experiment and from the models. 12 models for AG and 8 models for GA were built and screened using FPS. In this table, the computed values represent those closest to the mean distance from experiments. These values correspond to the ‘representative’ models shown in close up in Figure 3 part iii. For the computed distance values for all other models, see Supporting Materials Table S11. For TU, two proximity ratios obtained from experiments are shown that corresponds to two peaks (Figure 4A). Similarly, the proximity ratios obtained from the two models (Figure 4B) are shown.

| Calculated values (R or P) | Experiment |  | Model |  |
| --- | --- | --- | --- | --- |
| Labels: DNA | -93 | +94 | -93 | +94 |
| AG (R in Å) | 44.2 ± 0.84 | 50.0 ± 0.24 | 44.6 | 51.1 |
| GA (R in Å) | 49.0 ± 0.74 | 41.0 ± 0.74 | 50.2 | 40.6 |
| TU (P) | 0.54, 0.75 | 0.52, 0.74 | 0.55, 0.77 | 0.53, 0.72 |

We modelled these two nucleosomes taking the peak widths into consideration, thereby producing an ensemble of models to fit the major population peaks in the distance histograms (Figure S6). Figure 3A and B part ii (middle column) shows the ensembles and Table S11 in the Supporting Materials tabulate the distance values from the model ensembles. Column iii show close-up views of only the model from the ensemble whose computed distances match the experimental mean distance (Table S11 in the Supporting Materials). These models, representative of the mean distance, correspond to the 5NL0-type nucleosome for AG, and to the mode 7 frame 51 type, for the GA. The peak width of the distance histograms can reflect L-DNA dynamics, including A-tract-induced bending (Fig S6 in Supporting Materials for details) (30, 55–57). Thus, to sample L-DNA dynamics, as described in Materials and Methods (and Figure S6), structures of different arm-opening types of the nucleosomes were modeled. Slight orientational changes in the gH placements were also considered. The structural ensemble was screened using the FPS software (40) with the experimentally derived distances as input. A comparison of the experimentally derived mean distances and the model distances (Table 3 and Table S11 in the Supporting Materials) showed that the observed peak widths can indeed be explained by the dynamic fluctuations of the L-DNA.

The AG and GA models show that Zone B of the gH points towards the A-tract flank (Fig 3A and B, panel iii). Specifically, R42 (loop 1) and R94 (β2) are observed to be proximal to the minor groove of the A-tract, whereas the dye position (C77) is oriented away from the two L-DNAs, enabling free rotation of the donor dye.

A concern could be that the highly negatively charged Alexa 488 dye may tend to turn away from the DNA and affect the orientation of the gH, placing Zone B closer to the A-tract minor groove instead of Zone A. As reported by Zhou et al. (2020) in their recent cryo-EM structural studies (50), Zone A of the H1.0b subtype contains an arginine residue, R74, that can interact with the minor grooves of AT-rich DNA and could also be a suitable interaction candidate for our constructs. To investigate whether the presence of the dye on the α3-helix prevents such an interaction, we built an ensemble of alternate models (Supporting Materials Figure S14, Table S12) with R74 oriented towards the A-tract minor grooves of the AG and GA nucleosomes. This ensemble was built in the same way as the AG and GA ensembles, with four different arm-openings, and various poses of the gH. When the α3-helix points to the A-tract minor groove to enable R74 interactions, the dye attachment (T77C) site is still oriented away from both L-DNA arms, facing the gap between the L-DNAs, thus suggesting that the dye attachment site does not appear to influence or bias our result. The R74-A-tract ensemble was similarly screened using FPS (40). However, we observed that the computed distances between the A-tract and the gH (Table S9) are far higher (by 15 to 20 Å) than the experimentally derived mean distances, and beyond the distance ranges covered by the experimental peak widths of ±6 to ±7 Å around the mean distances. This analysis shows that, for AG and GA A-tract-containing nucleosomes, Zone B mediates the recognition of the A-tracts by the two arginines of the gH.

In all our models, the gH is positioned on-dyad. In this position, and with the modeled L-DNA separation, Zone B of the gH points towards the minor groove of the A-tracts. To investigate whether off-dyad binding might be consistent with our FRET data, we built off-dyad models for the 5NL0-type arm opening structures of the AG and GA constructs (See Supporting Materials figure S15, Table S13), while keeping the GVGA motif on the gH beta-hairpin loop (zone C) proximal to the major groove of the A-tracts. Using FPS, we calculated the possible inter-fluorophore distances. We found that an off-dyad positioning of the gH would result in inter-dye distances that are again much higher than, and inconsistent with, our experimental observations. Further support for an on-dyad positioning of the gH comes from the observations that LH subtype H1.0 associates with the nucleosome on-dyad (12, 50, 65). Comparison of the residues proximal to DNA in prior structural studies and the selected structures for our AG and GA models shows that the ‘FRET-restrained’ gH is located on the DNA in a similar manner to the previously observed on-dyad binding mode (Supporting Materials section S14). The close-up view of one of the models of the GA construct show that the beta-hairpin loop is positioned away from the dyad DNA. Although this structure corresponds to the mean distance, in the other conformers from the ensemble that represent the peak width, this interaction is recovered. Also, it can be noted that while the ‘closed’ conformation of the gH domain’s beta-hairpin loop in our GA model is not close enough to interact directly with the dyad DNA (Figure 3B iii), an ‘open’ conformation of the gH (66) could mediate such interactions in this pose. Both the open and closed conformations are observed in the crystal structure of chicken gH5 in isolation (66). Although never observed in conjunction with nucleosome in crystal structures, the ‘open’ conformation of the LH and the closed to open transition of an off-dyad bound gH5 has been observed in molecular dynamics simulations (32, 65), suggesting that this transition in the GA constructs could recover the beta-hairpin loop/NCP DNA interactions.

### The on-dyad bound H1.0b gH does not recognize thymines by their methyl group

It was proposed by Cui *et*.*al*. (2009) (31) that the affinity of gH to sequences with high AT content could be a result of hydrophobic interactions between the thymine methyl groups on the DNA and a hydrophobic patch on the gH, the GVGA motif in the beta-hairpin loop (zone C). This interaction was observed in molecular dynamics (MD) simulations of a nucleosome with a gH in the off-dyad binding mode (32), showing that the GVGA motif interacted with the thymine methyl groups exposed on the major groove of the DNA. Our LH, subtype H1.0b, contains the hydrophobic GVGA motif on the beta-hairpin loop (see Figure 1B i and ii). Why, then, was our gH unable to recognize the 64%-AT-rich DNA in MG and GM? The answer lies in the fact that the H1.0b subtype associates in an on-dyad fashion with the nucleosome. In our AG and GA models, we observed that the gH associated on-dyad and can access only the minor grooves of the L-DNA arms. To access the major groove of the L-DNAs, where the thymine methyl groups are exposed (denoted as blue spheres in the figures), the gH must associate in an off-dyad fashion. Thus, a gH positioned on-dyad is unable to employ the GVGA motif to recognize thymines but can recognize A-tracts due to DNA sequence-specific structural parameters, such as the well-documented narrowing of A-tract minor grooves. Arginines are well known to mediate such recognition (28, 34, 35, 67).

To clarify the potential contribution of thymine methyl groups, we made the TU construct, where both the plus and minus flanks have A-tracts. However, the minus flank was base-paired to dT while the plus flank was base-paired to deoxyuridine. The reason for using deoxyuridine was that dU lacks the C5 methyl group present in thymine. Thus, the TU minus flank is lined with methyl groups, whereas methyl groups are lacking in the plus flank. If the gH associated in an off-dyad binding mode, it could interact with the methyl groups on the minus L-DNA arm, and this would be expected to result in higher FRET with the A(T)-tract.

However, we observed that the proximity ratio distributions for the gH dye with the plus and minus dyes closely overlap (Fig. 4A, Table 2). The overlap shows that the gH, despite having the GVGA motif, has an equal preference for A-tracts paired to deoxyuridines lacking the methyl groups and to A-tracts paired to methyl group-containing thymines. This similarity suggests that the hydrophobic interaction between thymine methyl groups and the GVGA motif is not the only driving force governing the affinity of the gH for A-tracts. In addition, our observation that the gH does not show any preference towards random sequences having a higher AT-content (see above), further indicates that it is not just the hydrophobic interactions with the thymine methyl group, but other parameters of the DNA that come into play. By deconvoluting the proximity ratio profiles of the minus and the plus dyes of the TU construct, we observe two distinct populations within the peaks (proximity ratio 0.4 to 0.9). An additional low proximity ratio peak (mean proximity ratio = 0.2) could be a result of the LH being non-specifically bound to the nucleosome (Fig. 4A and Figure S10 and Table S7 in the Supporting Materials). The main peak (P : 0.4 to 0.9) can be fitted by two Gaussians, one at an intermediate mean proximity ratio of about 0.52 to 0.54 and one at a mean proximity ratio of 0.74 to 0.75, that appear in all the replicates in almost equal proportions. The peak widths of these two populations are comparable to the peak widths of the major populations in the other constructs (Table S7). These FRET profiles can be explained by the existence of two populations of the LH-nucleosome complexes, one with the gH dye closer to the AT flank, and the other with the gH dye closer to the AU flank. If there was only one position of the gH on the nucleosome and the two L-DNA arms of the TU construct bent in the same way, then we would have obtained only one population and would have observed a narrower peak.

**Figure 4:**
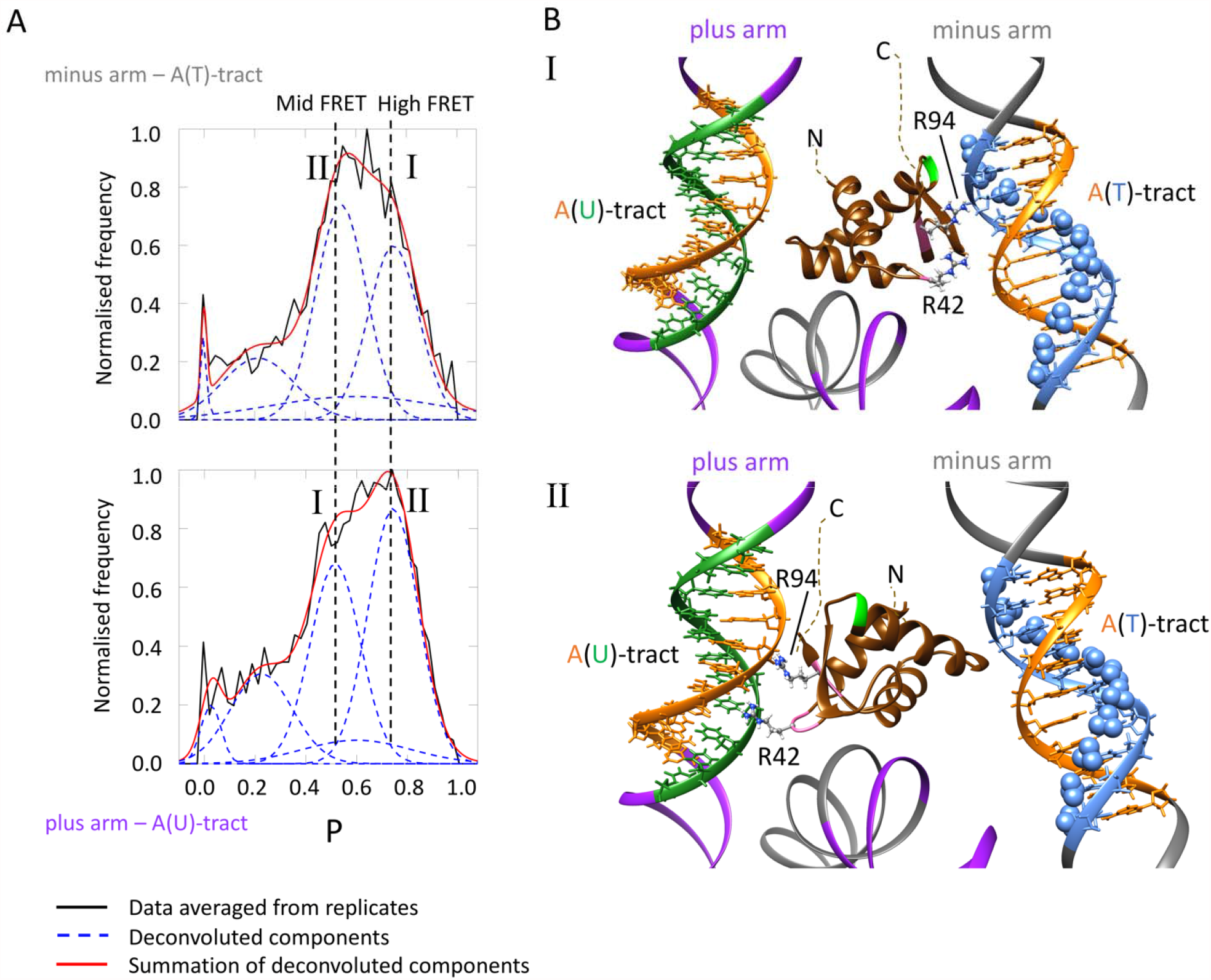
Evidence that the on-dyad bound LH H1.0b does not recognize A-tracts via the thymine methyl groups. **A**. Proximity ratio histograms obtained from the TU construct labelled on the minus or plus arms and on the gH (T77C site) of the full-length LH. The proximity ratio histograms between the minus arm and the gH completely overlap with the histogram obtained from the plus arm and the gH, showing that the gH has similar preferences for thymine- and uridine-containing A-tracts. Upon deconvolution of the histograms, the broad peak from P = 0.4 to P = 0.9 was fitted by two major populations. Population I (higher proximity ratio of gH with A(T)-tract minus arm, lower proximity ratio with A(U) tract plus arm) suggests that the gH is oriented towards the A(T)-tract minus arm. Population II (higher proximity ratio of gH with A(U)-tract plus arm, lower proximity ratio with A(T) tract minus arm) suggests that the gH is oriented towards the A(U)-tract plus arm. **B**. Models of the two populations: I has the gH oriented towards the A(T)-tract minus arm and II has the gH oriented towards the A(U)-tract plus arm. The flank sequences are colored orange (dA), light blue (dT) and green (dU). The thymine methyl groups are indicated by spheres. The computed proximity ratio from the models and the experimental mean proximity ratio of the I and II peaks are given in Table 3.

In each plot in Figure 4A, the high proximity ratio peak indicates that the gH is proximal to the corresponding labelled arm. The intermediate peak corresponds to the case where the gH is oriented away from the labelled arm and towards the other, unlabelled arm. The two populations are numbered I and II. We modelled structures representing these two populations and screened them in FPS (40) using the mean proximity ratio as the input. Since the peak widths of the two populations were comparable to the peak widths of the dominant populations in the AG and GA constructs (Table S6 and S7), we assume that the widths can be explained by similar L-DNA arm-dynamics. Population I (high FRET for minus arm and intermediate FRET for plus arm) has Zone B of the gH pointed towards the A(T)-tract on the minus arm and Population II (high FRET for plus arm and intermediate FRET for minus arm) has Zone B pointed towards the A(U)-tract on the plus arm. Table 3 shows the mean proximity ratio of the peaks and the computed FRET efficiencies of the two structures. They are comparable, suggesting that the two peaks indeed correspond to the two populations of nucleosomes with the gH oriented towards either the A(T)-tract or the A(U)-tract.

### The C-terminal domain of the *Xenopus laevis* H1.0b LH is not affected by the sequences of the L-DNA arms

To explore whether the CTD is affected by the flank sequences, we labelled the LH at residue 101. Fig. 5 shows the proximity ratio distribution between the donor dye on the CTD and the acceptor fluorophores on the plus and minus arms. We observed that for each of the nucleosomes, MG and GM (Fig. 5A and 5B, and Figure S11, Table S8 in the Supporting Materials for replicates), AG and GA (Fig. 5C and 5D, and Figure S12 for replicates), the proximity ratio distributions between the gH and the plus and minus arms nearly overlap with each other. The mean proximity ratios of the tail dye between the plus and minus arms have the maximum difference only in the AG construct (ΔP = 0.08) (Table 2 and Table S9). The tail dye in the AG construct is observed to have a higher proximity ratio for the GC-flank plus arm with respect to the A-tract flank minus arm. This suggests that the tail dye is oriented away from the flank towards which the gH is oriented, for the AG construct. This shift of the tail dye towards the arm that the gH domain is oriented away from is even lower (ΔP = 0.05, Table 2, Table S9) when the flank sequences are swapped in the GA construct.

**Figure 5:**
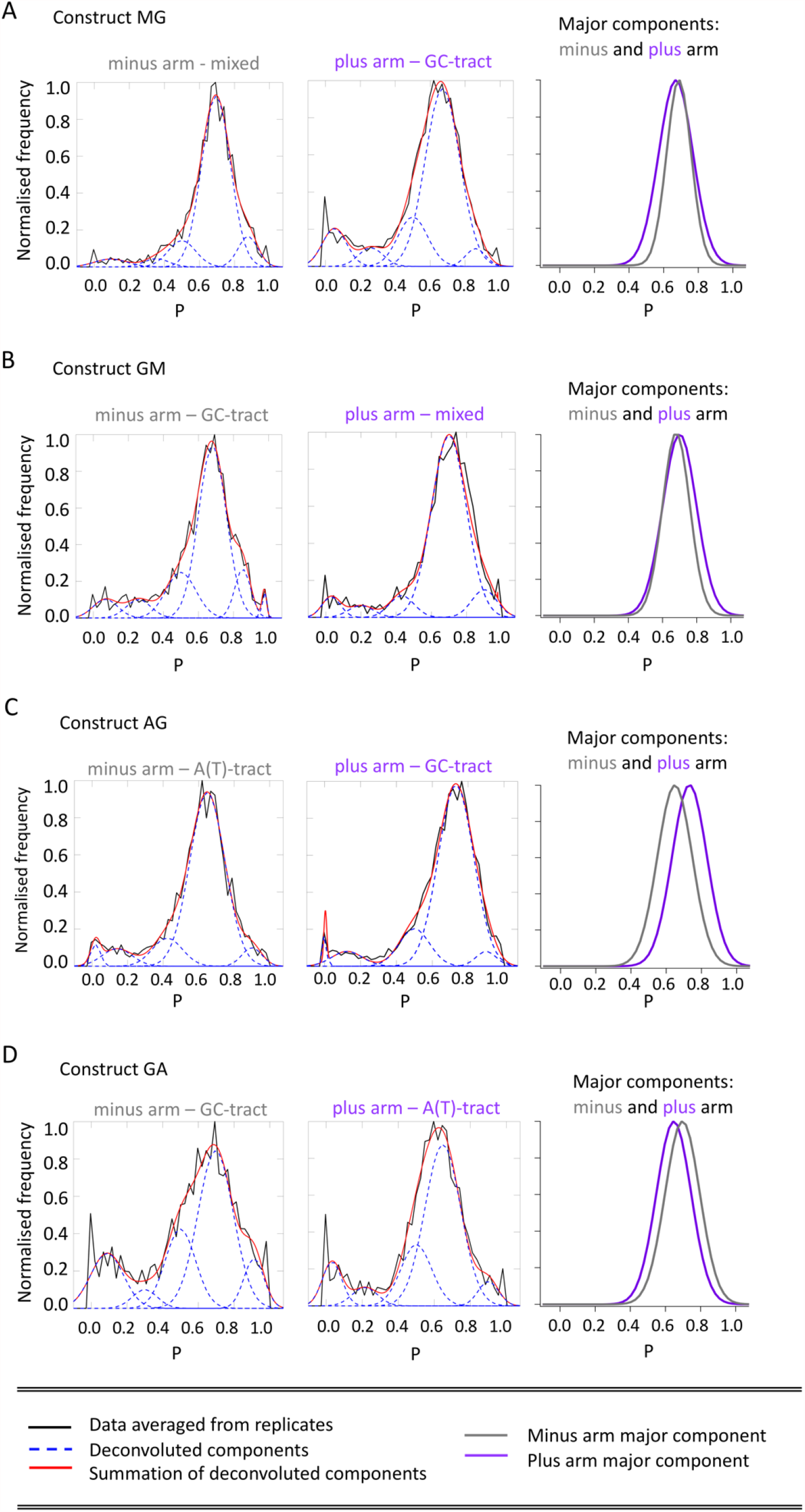
Proximity ratio histograms for nucleosomes MG (A), GM (B), AG (C), and GA (D) labelled on the plus (purple) or minus (grey) L-DNA arms and the CTD site of the LH (G101C). From left to right for each of the nucleosomes A-D, the plots show fitted proximity ratio histograms of the minus arm, the plus arm, and the major population peaks from the minus arm (grey) and the plus arm (purple) (height normalised) shown in one plot for comparison. The **(A)** MG construct and **(B)** GM construct show that the CTD-dye is nearly equidistant from both the minus and plus L-DNA arms. For **(C)** the AG construct, the CTD-dye is slightly closer to the GC-tract containing arm and **(D)** in the GA construct, the CTD-dye is again nearly equidistant from both the arms.

From our data for the tail dye, we cannot conclude whether the CTD is locally structured on the mononucleosome or it is disordered. But the good signal-to-noise ratio of the proximity ratio peak suggests that the dye is present on a more ordered part of the CTD. This could be because the labelling position at the 101st residue, is on the upstream end of the tail, just six residues away from the gH.

There is no effect of the examined flank sequences on the tail dye, in contrast to the observations for the gH dye. The tail dye is either equidistant from both the arms (MG, GM and GA constructs), or shifted slightly away from the flank that the gH is proximal to, as in the AG construct.

## Concluding discussions

The location of the LH on the nucleosome has garnered much attention because of its possible biological role: the two predominant modes by which the LH associates with the nucleosome (on- and off-dyad) have been suggested to have major implications for the compaction of higher-order structures of chromatin (8, 57). The positioning of the gH has been observed to be dependent on sequence variations (21) and post-translational modifications (13) of the gH.

However, despite numerous studies (24–27, 29, 31, 32) showing the preference of the LH for AT-rich regions, the contribution of the DNA sequence to the positioning of the LH is unclear. In the present work, we examined the possible effect of the DNA sequence on the location and the orientation of the LH. We first tested the preference of the LH for a 64% AT-rich mixed sequence vs. a purely GC-tract, in the MG and GM constructs. Surprisingly, in the MG construct, the gH dye showed higher FRET efficiency with the L-DNA arm containing a purely GC sequence than with the 64% AT-rich mixed sequence. In the GM control construct, the gH showed similar proximity ratio profiles for both the L-DNA arms, showing no preference for either the purely GC sequence or the 64% AT-rich sequence. These apparently contradicting observations led us to question possibilities such as the asymmetry of the non-palindromic Widom 601 sequence, the variable unwrapping of the L-DNAs, or the proximity of the pure-GC L-DNA to the gH in the MG construct. By using suitable controls with the flank sequences swapped (GM) we rule out these possibilities. We suggest that the apparent preference of the gH for the pure-GC arm in the MG construct could be due to the orientation of the gH.

When we substituted the 64% AT-rich mixed sequence with an A-tract (AG and GA constructs), we consistently observed that the gH dye showed a high proximity ratio for the A-tract containing L-DNA.

Although our gH showed a clear proximity towards the A-tract flank in the constructs AG and GA, the CTD dye did not show any DNA-sequence preference. It was equidistant from both the L-DNAs in the constructs MG and GM and was oriented away from the A-tract in the AG and GA constructs (Figure 5). From our results with the CTD dye, we cannot conclude how the rest of the CTD is associated with the L-DNA arm. The CTD had been observed to associate with one or both L-DNA arms, depending on the salt concentration, in coarse-grained simulation studies (16), and with one L-DNA arm in cryo-EM studies when a palindromic Widom 601 sequence was used (12). But recent cryo-EM work from Zhou *et al*. (2021) (23) using the same DNA sequence as our MG construct (non-palindromic Widom 601), showed that the CTD of the H1.0 subtype associate similarly with both the L-DNA arms. Although recognition of AT-rich DNA has been attributed to the CTD in the work of Roque *et al*. (2004) (29), in our study we see that the labeling position on the CTD is oriented away from the A-tract. Thus, from our studies, we suggest that recognition of AT-rich DNA is imparted largely by the gH domain.

Our distance-restrained models showed that the gH is located on-dyad and is associated in a canonical fashion. Out of the three DNA proximal zones, Zone B was observed to be close to the minor groove of the A-tract (minus arm of AG, plus arm of GA). In this zone, two amino-acid residues likely mediate the A-tract recognition. A highly conserved arginine (R42) is present in the loop 1 region of Zone B. A second arginine (R94), conserved in H1.0 isoforms (see Supporting Materials Table S15), is present on the beta-sheet at the C-terminal end of the gH. These two arginines protrude into the characteristically narrow minor groove of the A-tract. A-tract regions, which are abundantly present in SAR (scaffold attachment regions) (68) and satellite sequences (27), are known to have the narrowest minor groove, which has a strong negative electrostatic potential. Arginines, being highly positively charged, interact electrostatically with the minor grooves of A-tracts (28, 34, 35), and are preferred over lysines in this particular interaction (67) because of the guanidium group that makes strong hydrogen bonds with the DNA (13) and has a lower electrostatic desolvation penalty compared to the smaller amino group of lysine (69). Electrophoretic mobility shift assays between lysine-rich somatic LH and arginine-rich sperm LH associated with DNA (70) showed a greater DNA-binding capability of arginine-rich sperm LH. This was also reflected in binding assays (29) that showed that arginine-rich protamine nuclear proteins were selectively bound to A-tract rich DNA. Based on these findings and our observations, we propose that, specifically in H1.0 isoforms, the Zone B region, with its two arginines in a ‘pincer’-like motif, is a key contributor to A-tract recognition by LHs. Functionally, this arginine ‘pincer’ motif is similar to the well-known AT-hook motif known to mediate recognition of AT-rich DNA tracts (71). However, structurally, the AT-hook motif consists of two arginines close to each other in the sequence (Pro-Arg-Gly-Arg-Pro) (71) whereas in the ‘pincer-like’ motif of the H1.0b Zone B, the two arginines, 42 and 94, are close in space. Mutational studies on these two conserved arginine residues had been performed in vitro by Goytisolo *et al*. (1996) (72) and Duggan *et al*. (2000) (73*), in vivo* by Brown *et al*. (2006) (18), and *in silico* by Öztürk *et al*. (20). All these studies showed that mutating these arginines to glutamate (72) or alanine (18, 20, 73) decreased DNA binding by the LH. However, the effects of these mutations on binding AT-rich DNA have not been studied. In future work, it would be interesting to investigate how mutating the arginines to other residues, such as lysine, affects the A-tract recognition.

The mode of recognition of AT-rich DNA that we propose, differs from that proposed by Cui and Zhurkin (2009) (31) and observed in simulations by Öztürk *et al*. (2016) (32). Both the groups suggested that the gH, when positioned off-dyad, can directly mediate hydrophobic interactions with thymine methyl groups exposed on the major groove of the L-DNA arm, and that this interaction is mediated by the hydrophobic GVGA motif on the beta-hairpin loop (Zone C, Figure 1B). The LH that we use for our experiments, H1.0b, contains the GVGA motif. It has also been observed to associate in an on-dyad fashion, both in our models and in previous structure determination studies (12, 50) and in a computational study (65), where it was shown that the on-dyad binding mode of the H1.0 subtype is enthalpically favoured. We suggest that the mode of association of the LH isoform, i.e. on-dyad, could be the reason why our H1.0b LH, despite containing the GVGA motif, is unable to recognize the 64% AT-rich mixed flank in the MG and GM constructs. To access the major groove, and thereby interact with the thymine methyl groups of the 64% AT-rich region through hydrophobic interactions, the gH has to associate in an off-dyad binding mode (31, 32). To further prove that our LH does not mediate A-tract recognition via interactions with the thymine methyl group, we constructed the TU nucleosome containing A-tracts complementary to thymine as well as uridine. A-tracts complementary to deoxyuridines have very similar minor groove widths to A-tracts complementary to thymines. Replacing thymines with uridines has previously been observed not to confer any significant change in the minor groove width (74). The proximity ratio profiles from the TU construct show the presence of two populations, one with the gH domain oriented towards the thymine-tract minor groove, and the other with the gH domain oriented towards the deoxyuridine-tract minor groove.

Considering previous studies on AT-rich DNA recognition by the LH (31, 32) and the results reported here, as well as very recent cryo-EM structures (50) showing that R74 on the α3-helix could mediate recognition of AT-rich DNA via minor groove interactions, we suggest that there is more than one mechanism by which LHs can recognize AT-rich sequences.

Taken together, we propose that all three DNA-interacting zones on the gH domain, A, B, and C, can recognize AT-rich DNA (Figure 6). Zone A has arginines dotting the α3 helix. In Zone B, R42 on loop 1 is highly conserved (see Table S15 in the Supporting Material for Zone A, B, and C sequences amongst LH isoforms across organisms). These two zones are proximal to L-DNA minor grooves in an on-dyad positioned gH domain and, as shown in this work, can identify A-tracts indirectly due to the sequence-induced structural changes in the DNA, such as the narrowing of the minor groove width in A-tracts. The hydrophobic Zone C, although less well conserved, can mediate direct recognition of AT-rich DNA, via major groove interactions, when the LH associates in an off-dyad binding mode. These multiple means of recognizing AT-rich DNA and A-tracts may explain why all LH isoforms have been observed to bind to and compact A-tract-containing SAR regions and AT-rich constitutive heterochromatin (75, 76). Finally, the present work indicates directions for future studies to comprehensively characterize the different mechanisms of nucleosomal positioning and orientation of LHs and their dependence on DNA and LH sequences, post-translational modifications, as well as the possible competitive interactions of core histone tails. Our study shows a new mode of recognition of A-tracts by the LH, and a DNA sequence-dependent variation in the positioning and orientation of the LH. To study the functional implications of this LH binding mode, chromatin arrays constructed with A-tracts bounding the NCP regions of select nucleosomes could be investigated.

**Figure 6:**
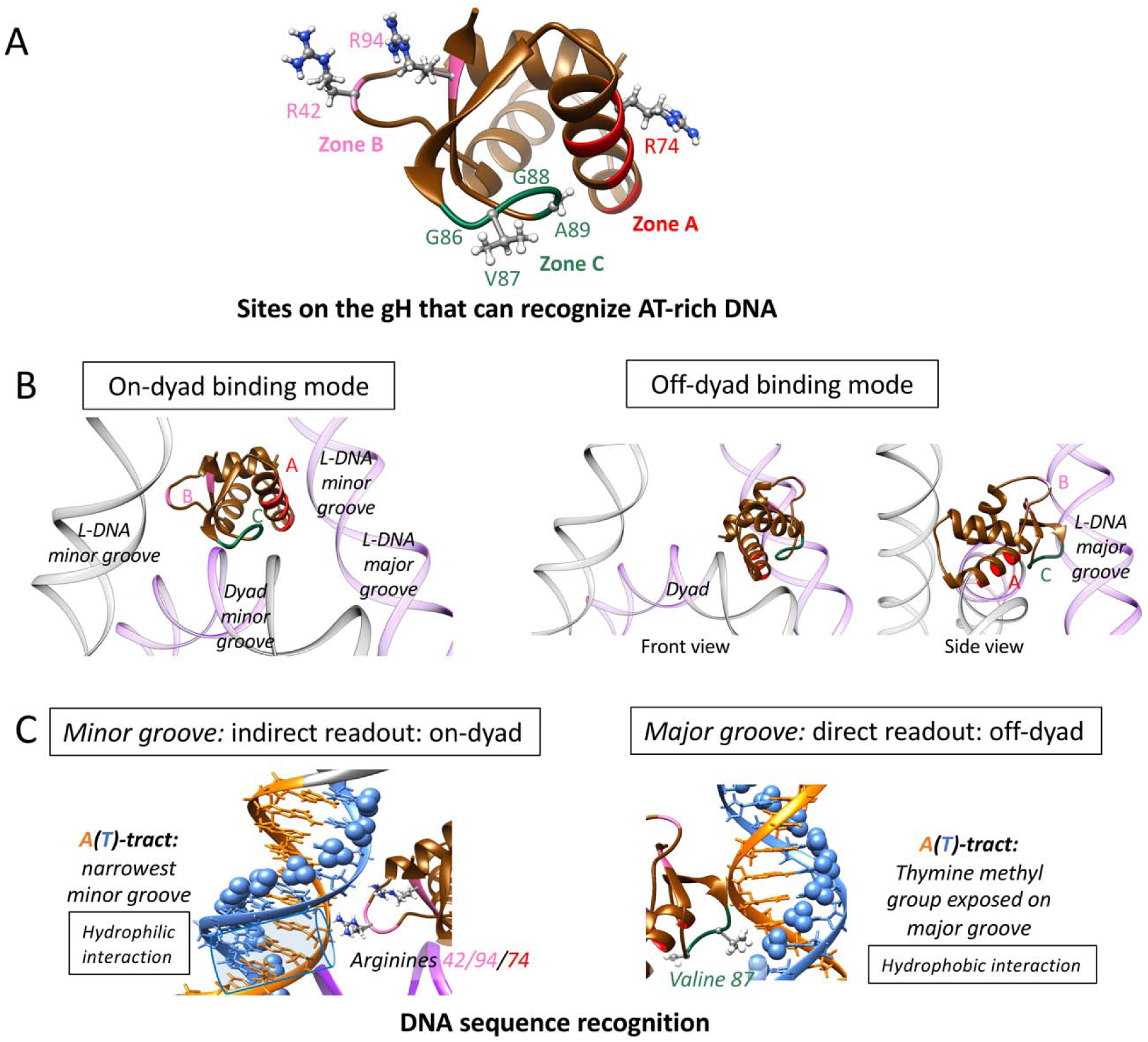
Schematic diagram to show the different mechanisms by which the globular domain of a linker histone recognizes AT-rich DNA. **A**. Residues suggested to mediate recognition of AT-rich DNA, on the three DNA-interacting zones of the gH domain are indicated. **B**. Representative on-dyad - (9, 12, 21, 23) and off-dyad (21, 31, 32, 65) binding modes showing how the different DNA-interacting zones of the gH are arranged with respect to the linker and nucleosomal DNA. C. The mechanism of recognition of A-tracts or AT-rich DNA by the gH when bound on-dyad (revealed in this work) or off-dyad (add refs).

## Abbreviations

LH: linker histone
L-DNA: linker DNA
gH: globular domain
NCP: nucleosome core particle
FRET: fluorescence/Förster resonance energy transfer
ALEX: Alternating Laser Excitation
FPS: FRET Positioning and Screening

## Data and software availability

Data for this work are available at https://doi.org/10.5281/zenodo.4686909. The software used to obtain proximity ratio and stoichiometry histograms, AlexEval, is available at: https://github.com/sisbaner/AlexEval.

## Author contributions

M.D., K.T., and R.C.W. formulated the project. M.D. and K.T. designed the experiments. M.D., M.A.O and R.C.W. designed the modelling workflow. M.D. performed the research, analyzed the data, and wrote the manuscript with suggestions from M.A.O., K.T., and R.C.W. S.I. developed the AlexEval software. All authors contributed to the finalization of the manuscript.

## Acknowledgements

We dedicate this work to the memory of the late Prof. Dr. Jörg Langowski, a remarkable pioneer in the field of chromatin biophysics. We are very grateful to Prof. Jeffrey Hayes (University of Rochester Medical Center) and Dr. Amber Cutter for kindly providing us with the plasmids and protein for unlabelled and labelled LH. We thank Nathalie Schwarz for DNA purification and, together with Maria Mildenberger and Gabriele Mueller, for protein purification. M.D. is thankful to Angga Prawira for measuring some technical replicates of AG and MG nucleosomes. We are grateful to Dr. Kathrin Lehmann for assistance in designing the MG nucleosome and for helpful feedback on the manuscript.

