## Supporting Materials for "DNA sequence-dependent positioning of the linker histone in a nucleosome: a single-pair FRET study"

**Materials – Sequences and labelling positions**

**DNA sequences:** All sequences are 226 bp long and run from 5’ to 3’. Only ‘forward’ strands are shown for AG, GA, MG, and GM constructs. For TU constructs, both forward and reverse strands are shown. The core DNA (shown in bold) has the non-palindromic Widom 601 sequence. The original Widom 601 sequence is modified at the flank sequences (underlined) in constructs AG, GA, TU, and GM (see Table 1, Results section). The dyad base (italics) is numbered 0. Bases to the left of the dyad (0) shown in grey are assigned negative values and bases to the right of the dyad shown in purple are assigned positive values. This colour scheme is consistent in all the proximity ratio plots. The DNAs were labelled on thymines with Alexa 594 (acceptor dye). In nucleosomes with the donor dye on the LH, the DNA was labelled on the -93 T (minus-arm label) on the forward strand or on +94 T on the reverse strand (denoted by **A^(+94)^** on the forward strand sequence given here). In AG, GA, and TU DNA, the **[ ]** brackets enclose the 193 bp of DNA that were used in modelling.

**AG:**

ATCCGACTGGCACCGG [ CAA**T^(-93)^[Alexa 594]** GTCGCTGTTCCAAAAAAAAAA**AGGATGTATATATCTGACACGTGCCTGGAGACTAGGGAGTAATCCCCTTGGCGGTTAAAACGCGGGGGACA*G*^(0)^CGCGTACGTGCGTTTAAGCGGTGCTAGAGCTGTCTACGACCAATTGAGCGGCCTCGGCACCGGGATTCTCCAG**GGCGGCCGCGTATAGGGTCC**A^(+94)^ [Alexa 594]** TC ] ACATAAGGGATGAACTC

**GA:**

ATCCGACTGGCACCGG [ CAA**T^(-93)^ [Alexa 594]** GTCGCTGTTTACGCGGCCGCC**CTGATGTATATATCTGACACGTGCCTGGAGACTAGGGAGTAATCCCCTTGGCGGTTAAAACGCGGGGGACA*G*^(0)^CGCGTACGTGCGTTTAAGCGGTGCTAGAGCTGTCTACGACCAATTGAGCGGCCTCGGCACCGGGATTCTCCCT**TTTTTTTTTTGGTAGGGTCC**A^(+94)^ [Alexa 594]** TC ] ACATAAGGGATGAACTC

**TU:** In the complementary strand, the + flank has dU and the – flank has dT. The dU-tract was incorporated into the reverse primer. The strands were designed to have the same bases on the same strands (i.e. A-tract on minus and plus sides of forward strand, T- and U-tract on the reverse strand) to prevent the possibility of a hairpin loop formation, due to intra-strand hybridisation of the A-tract with the T- or U-tract.

Forward strand:

ATCCGACTGGCACCGG [ CAA**T^(-93)^ [Alexa 594]** GTCGCTGTTCCAAAAAAAAAA**AGGATGTATATATCTGACACGTGCCTGGAGACTAGGGAGTAATCCCCTTGGCGGTTAAAACGCGGGGGACA*G*^(0)^CGCGTACGTGCGTTTAAGCGGTGCTAGAGCTGTCTACGACCAATTGAGCGGCCTCGGCACCGGGATTCTCCCA**AAAAAAAAAAGGTAGGGTCCATC ] ACATAAGGGATGAACTC

Reverse strand: from 5’ to 3’. Instead of thymines in the plus (purple) strand, there are deoxyuridines (U).

GAGTTCATCCCTTATGT [ GA**T^(+94)^ [Alexa 594]** GGACCCTACCUUUUUUUUUU**UGGGAGAATCCCGGTGCCGAGGCCGCTCAATTGGTCGTAGACAGCTCTAGCACCGCTTAAACGCACGTACGCG*C*^(0)^TGTCCCCCGCGTTTTAACCGCCAAGGGGATTACTCCCTAGTCTCCAGGCACGTGTCAGATATATACATCCT**TTTTTTTTTTGGAACAGCGACATTG ] CCGGTGCCAGTCGGAT

**MG:**

ATCCGACTGGCACCGGCAA**T^(-93)^ [Alexa 594]** GTCGCTGTTCAATACATGCAC**AGGATGTATATATCTGACACGTGCCTGGAGACTAGGGAGTAATCCCCTTGGCGGTTAAAACGCGGGGGACA*G*^(0)^CGCGTACGTGCGTTTAAGCGGTGCTAGAGCTGTCTACGACCAATTGAGCGGCCTCGGCACCGGGATTCTCCAG**GGCGGCCGCGTATAGGGTCC**A^(+94)^ [Alexa 594]** TCACATAAGGGATGAACTC

**GM:**

ATCCGACTGGCACCGGCAA**T^(-93)^[Alexa 594]** GTCGCTGTTCGCGGCCGCCAC**AGGATGTATATATCTGACACGTGCCTGGAGACTAGGGAGTAATCCCCTTGGCGGTTAAAACGCGGGGGACA*G*^(0)^CGCGTACGTGCGTTTAAGCGGTGCTAGAGCTGTCTACGACCAATTGAGCGGCCTCGGCACCGGGATTCTCCAG**GCATGTATTGTATAGGGTCC**A^(+94)^ [Alexa 594]** TCACATAAGGGATGAACTC

**Linker histone sequence:**

Sequence of the full-length H1.0b (*Xenopus laevis*). The underlined portion is the globular domain (gH) used for modelling. The LH was labelled with Alexa 488 (donor dye) either on the gH (at residue 77 with threonine mutated to cysteine) or on the C-terminal tail (at residue 101 with glycine 101 mutated to cysteine).

MAENSAATPAAKPKRSKALKKSTDHPKYSDMILAAVQAEKSRSGSSRQSIQKYIKNHYKVGENADSQIKLSIKRLV**C^77^[Alexa 488]** SGALKQTKGVGASGSFRLAKADE**C^101^[Alexa 488]** KKPAKKPKKEIKKAVSPKKVAKPKKAAKSPAKAKKPKVAEKKVKKVAKKKPAPSPKKAKKTKTVKAKPVRATKVKKAKPSKPKAKASPKKSGRKK

**Materials – Gels: protein labelling**

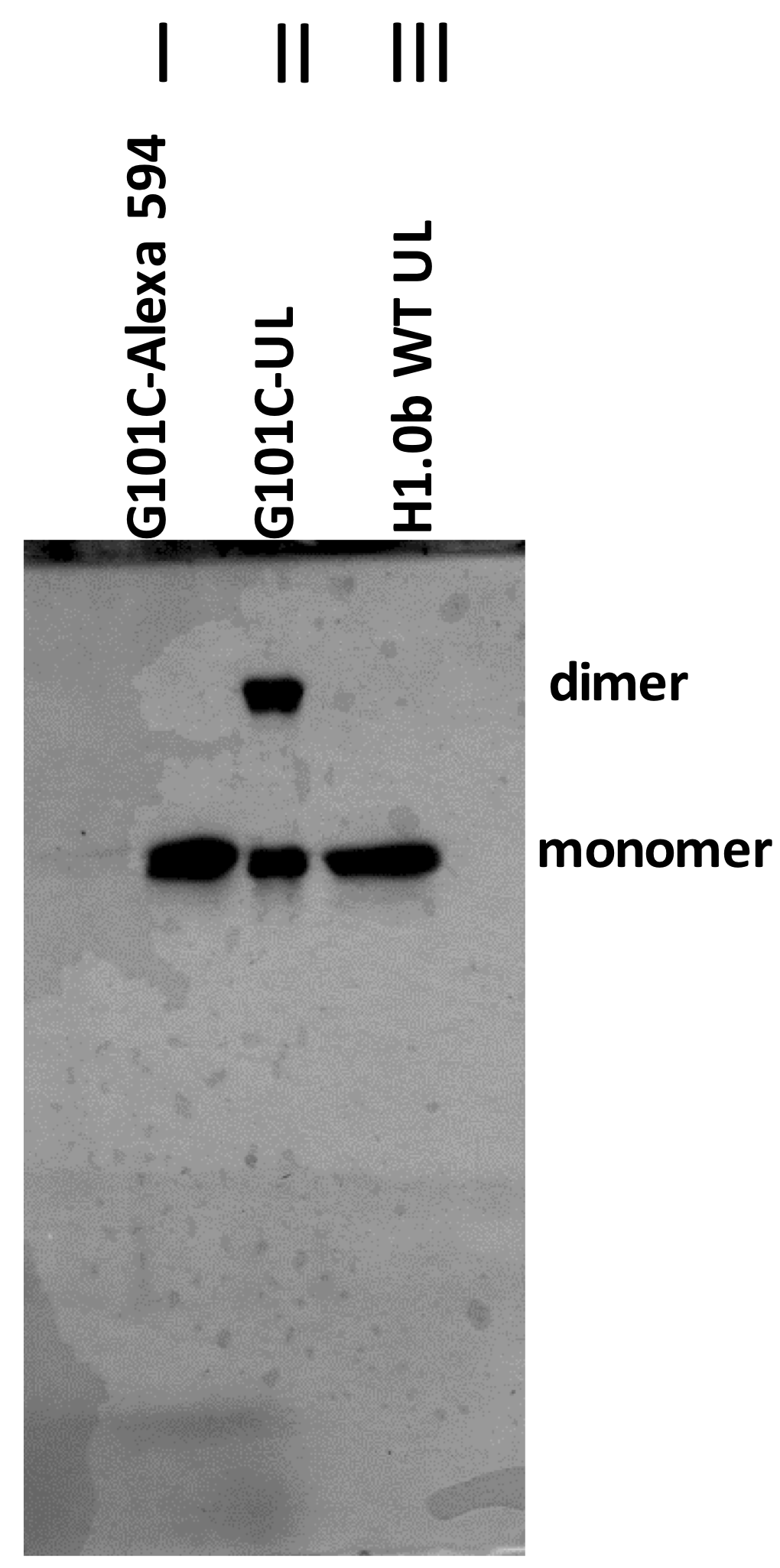
**Fig S1:** Denaturing gel electrophoresis of CTD-labelled and unlabelled linker histone. 15% SDS-PAGE (Coomassie Blue stained) of wild-type (lane III), mutant (lane II), and labelled mutant LH H1.0b (lane I), run at 150V (21.4 V/cm) for 1 hour. Wild-type H1.0b (lane III) shows no homodimer formation, because it does not contain any cysteine residues. The G101C unlabelled mutant (lane II) shows the presence of a high MW fraction, suggesting dimer formation. Labelling the G101C mutant (lane I) at the cysteine prevents further dimerization.

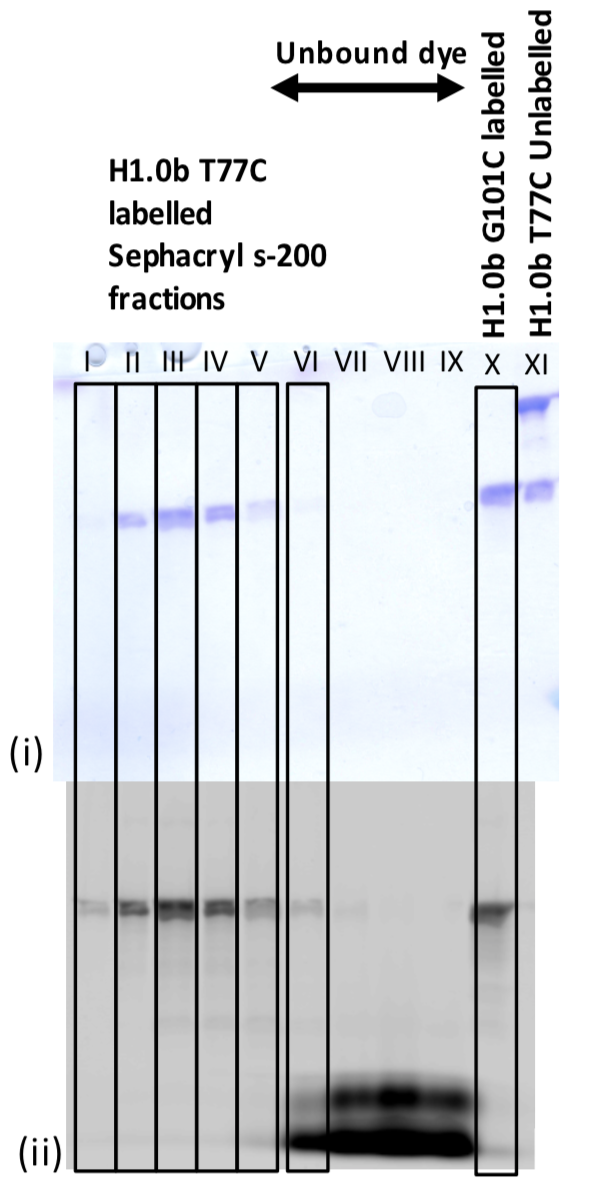
**Fig S2:** Denaturing gel electrophoresis of gH-labelled and unlabelled linker histone. (i) Coomassie Blue stained 15% SDS-PAGE, run at 150V (21.4 V/cm) for 40 minutes. (ii) same gel before staining, imaged by a Typhoon 9400 scanner (1) (GE Healthcare) with excitation at 488 nm, emission between 500 and 540 nm, showing Alexa 488 labelled proteins.

Lanes I to VI: Labelled T77C mutant LH that was passed through Sephacryl S-200 to remove unbound dye. Unbound dyes show up at the base of the gel (lanes VII to IX) in the Typhoon image (ii) but not in the Coomassie stained gel (i). In image (i), H1.0b T77C unlabelled (lane XI) shows up as a high MW dimer (top band) (due to the presence of cysteine) and a monomer. The labelled G101C mutant (lane X), shown for comparison, is a monomer, as is the labelled T77C mutant LH (lanes I to VI).

**Materials – Nucleosome gels:**

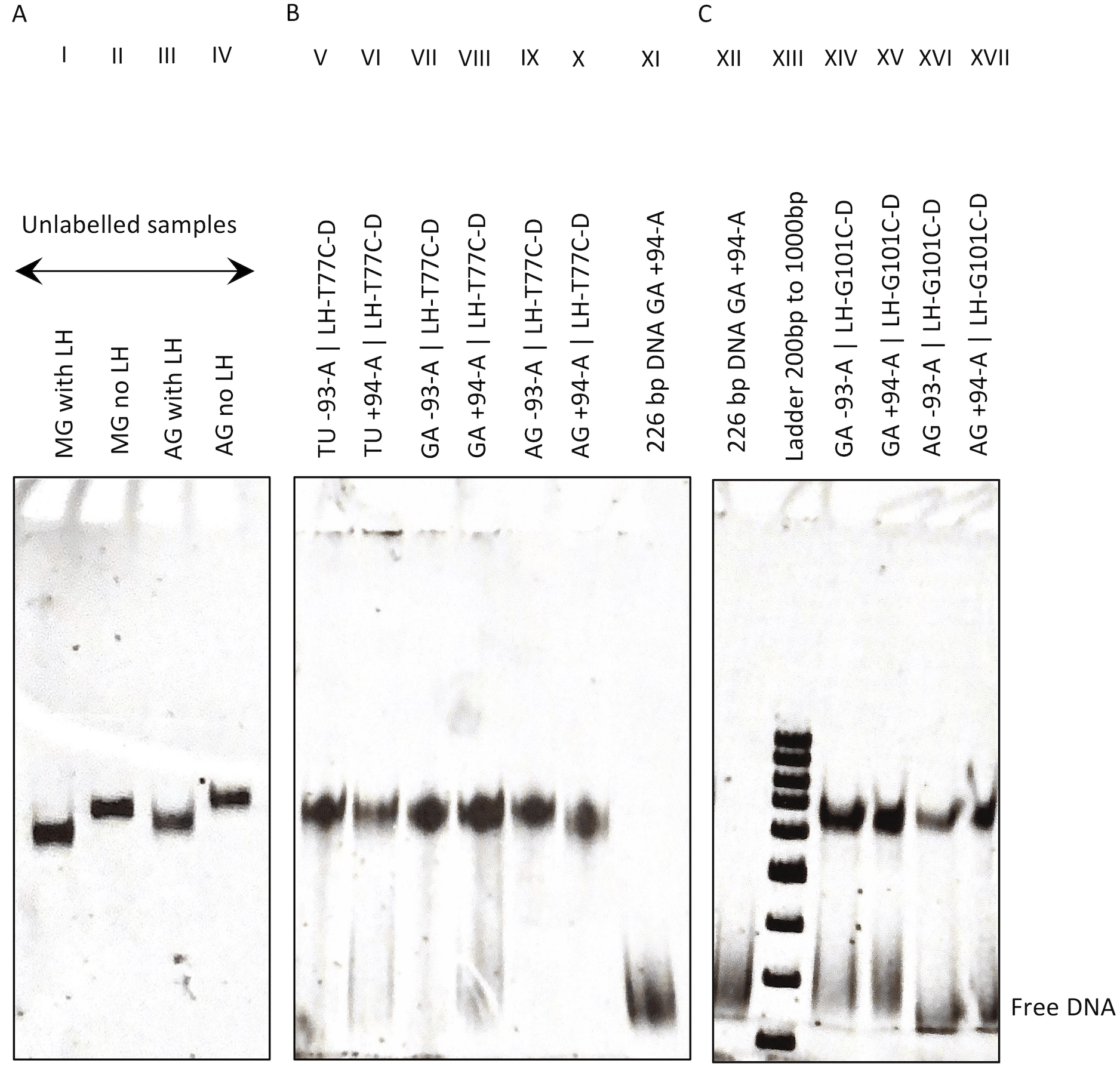

**Fig S3A-C:** Three 6% native (non-denaturing) polyacrylamide gel electrophoreses were performed to check for the presence of reconstituted nucleosomes (see Materials and Methods). All three gels were run at 70V (10V/cm), for 1.5 hours, EtBr stained. 50 ng of samples/well were used.

**Gel A** shows as a control, two samples without the LH (lane II: MG nucleosome without LH and lane IV: AG nucleosome without LH). Despite higher MW of the LH-containing nucleosomes (lanes I and III), the higher compaction of these nucleosomes results in their faster migration through the gel pores, compared to the lower MW but less compact, LH-lacking nucleosomes (lanes II and IV).

**Gel B:** Shown are nucleosomes with gH labelled LHs and labelled L-DNA, TU (lanes V, VI), GA (lanes VII, VIII) and AG (lanes IX, X). Lane XI shows a 226 bp DNA (GA construct) labelled at the +94^th^ base.

**Gel C:** Lane XII shows a 226 bp GA DNA labelled at +94. Lane XIII shows a 100 bp ladder marker (bottom to top: 200 to 1000bp). Shown are the GA (XIV and XV) and AG (XVI and XVII) nucleosomes with CTD-labelled LH. Some samples show the presence of free DNA (lanes VI, VIII, XIV to XVII).

**Methods – Obtaining inter-dye distances from single-molecule FRET and bulk spectroscopy**

The FRET efficiency is inversely related to the distance between the donor and acceptor fluorophores as given by the equation:

$$E= \frac{R_{0}^{6}}{R_{0}^{6}+R^{6}}$$

[1]

R_0_ is the Förster distance between the fluorophore pairs, and E is the FRET efficiency. Extracting R or inter-fluorophore distances from FRET efficiency is a multi-step process that involves both bulk spectroscopic measurements and single-molecule (alternating laser excitation-FRET) ALEX-FRET (2, 3), see Figures S4 and S5.

First, to calculate the Förster distance (in Å), we used the following formula:

$$R_{0}={0.211\left( \kappa^{2}.n^{-4}.Q_{D}.J\left( \lambda\right) \right)}^{1/6}$$

[2]

where κ^2^ is a dipole orientation factor and n is the refractive index (4). Q_D_ is the quantum yield of the donor fluorophore (Alexa 488) in ‘donor-only’ labelled nucleosomes. Secondly, the overlap integral J(λ) was calculated from the donor-only and acceptor-only (Alexa 594) equivalents of the nucleosome construct studied (5) using the following equation :

$$J\left( \lambda\right)= \frac{\int_{0}^{\infty} F_{D}\left( \lambda\right).\varepsilon_{A}\left( \lambda\right). \lambda^{4}d\lambda}{\int_{0}^{\infty} F_{D}\left( \lambda\right)d\lambda}$$

[3]

Here F_D_ is the fluorescent intensity of the donor dye in donor-only nucleosome, ε_A_ is the molar extinction coefficient of the acceptor in the acceptor-only nucleosome.

Absorbance measurements in bulk, at UV-visible range (220 to 750 nm) were carried out using a Cary 4E spectrometer (Varian, Mulgrave, Australia). For fluorescence measurements in bulk, a SLM-AMINCO 8100 fluorescence spectrometer (SLM, Urbana, IL) was used and emission spectra were collected in the wavelength range between 500 and 750 nm (6).

For calculating quantum yields of donor-only nucleosomes containing Alexa 488 or donor dye, fluorescein isothiocyanate in 0.1M NaOH was used as the reference. The following formula was used to calculate the quantum yields of Alexa 488 in nucleosomes:

$Q_{D}=Q_{\mathrm{ref}}\frac{{n_{D}}^{2}}{{n_{\mathrm{ref}}}^{2}} \frac{F_{D}}{\mathrm{OD}_{D}} \frac{\mathrm{OD}_{\mathrm{ref}}}{F_{\mathrm{ref}}}$	[4]

Q_ref_ refers to the quantum yield of fluorescein isothiocyanate in 0.1M NaOH: 0.95,

n_D_ is the refractive index of the sample: 1.4 (4), and n_ref_ is the refractive index of 0.1M NaOH: 1.33 (7). F_D_ and OD_D_ refer to the integrated fluorescence intensity and the absorbance for Alexa 488 (donor or *D* dye). F_ref_ and OD_ref_ refer to the integrated fluorescence intensity and absorbance of the reference.

From single-molecule FRET experiments, we obtain the proximity ratio. The proximity ratio is the number of photons detected in the acceptor channel, N_A_, divided by the total number of detected photons in the two channels.

$$P= \frac{N_{A}}{N_{D}+N_{A}}$$

[5]

The proximity ratio is related to the FRET efficiency by an instrumental detection factor, γ. We calculate the γ value using the method described by Lee et al. (2005) (2) and Kapanidis et al. (2004) (3) by measuring the proximity ratio and stoichiometry of two standard FRET samples: 30 bp B-DNA oligonucleotides, having FRET labels 10 and 21 bp apart. We assumed the quantum yields of the dyes in our FRET standards to be similar to those for our nucleosome samples. FRET efficiency E and proximity ratio P are related by the following equation:

$$E= \frac{P}{\gamma-P(\gamma-1)}$$

[6]

From P and γ, we calculate E. Incorporating E and R_0_ into equation 1, we obtain the distances, R between the fluorophores.

For modelling, we used the software FRET Positioning and Screening (FPS) (8) to screen for structures showing computed interfluorophore distances similar to R.

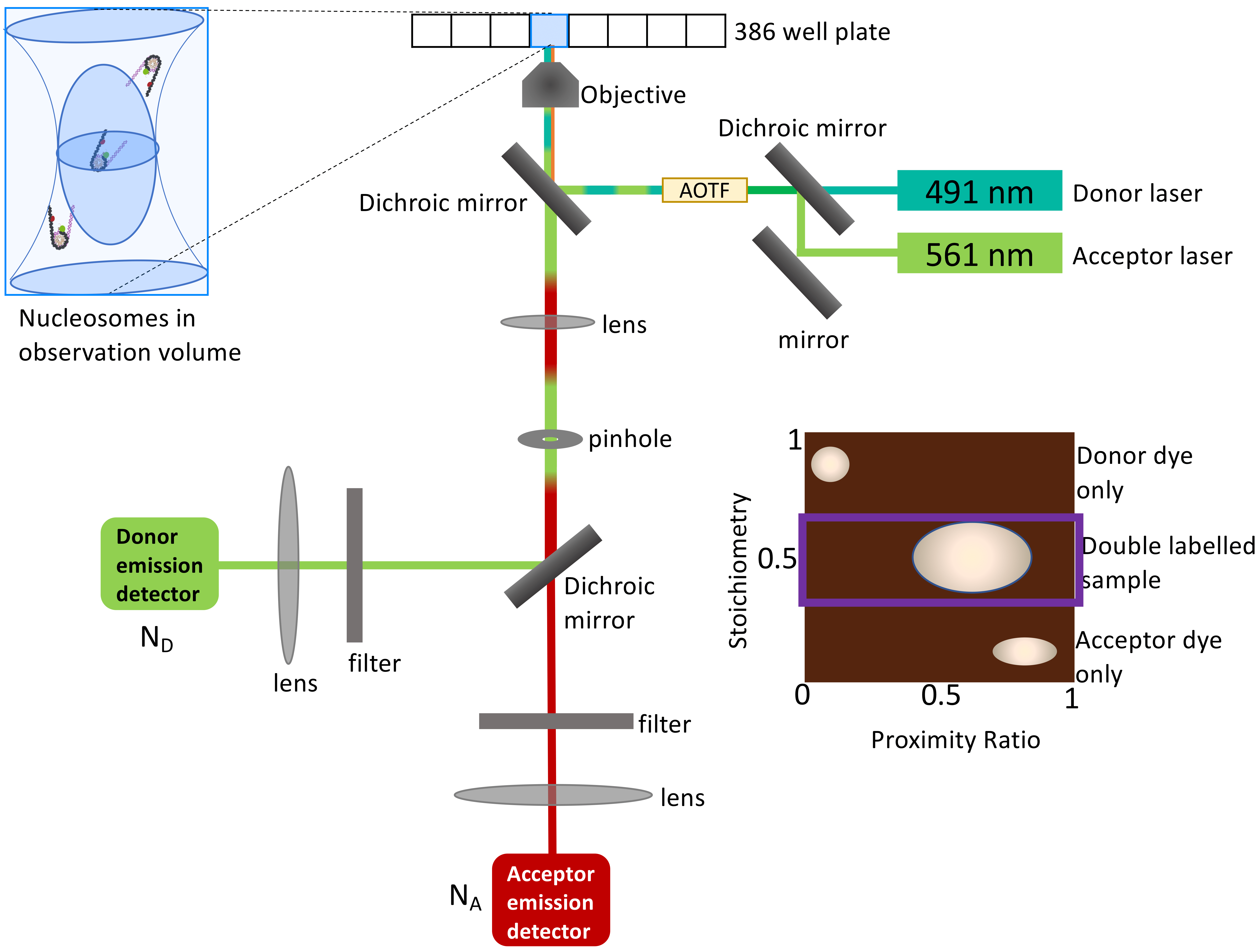

**Fig S4:** Schematic diagram of the in-house ALEX-spFRET setup. Inset: Schematic plot of stoichiometry vs. proximity ratio obtained from ALEX. The part of this plot, bounded by the purple rectangle, having a stoichiometry of 0.5, denotes the doubly labelled sample.

**Table S1 – Bulk spectroscopic values for deriving Förster distance:** We assume the refractive index of the buffer to be 1.4 (4). For the donor only sample, we used an AG nucleosome labelled only at LH-T77C (globular domain label) with Alexa 488 dye. The quantum yield of the donor-only sample (measured for a nucleosome comprising of unlabelled DNA, and LH labelled only on T77C in the gH) was 88%. Based on the measured values of the anisotropies of the donor and acceptor fluorophores of the doubly labelled (dl) samples (rD and rA), we assumed the value of κ^2^ to be 2/3. For each sample, values were calculated for a single replicate.

| **Constructs** | **Dye**  **specifications** | **QY_D_ (Alexa 488 (D)-only sample)** | **rD (dl sample)** | **rA (dl sample)** | **J(λ)**  (cm^-1^M^-1^nm^4^) | **κ^2^**  assumed | **Refractive index** | **R_0_ (Å)** |
| --- | --- | --- | --- | --- | --- | --- | --- | --- |
| Protein+DNA  **AG**  LH-T77C-Alexa 488 /  DNA +94-Alexa 594 | Alexa 488–C5-maleimide (protein)  Alexa 594–C6-dT (DNA) | 0.88 | 0.19 | 0.19 | 1.46x10^15^ | 2/3 | 1.4 | 52 |
| Protein+DNA  **GA**  LH-T77C-Alexa 488 /  DNA +94-Alexa 594 | Alexa 488–C5-maleimide (protein)  Alexa 594–C6-dT (DNA) | 0.88 | 0.22 | 0.11 | 1.45x10^15^ | 2/3 | 1.4 | 52 |

**Table S2 - Förster distances for Alexa 488-Alexa 594 dye pairs from the literature:** We compare our calculated R_0_ value to those obtained from the literature, by converting the R_0_ values obtained from literature with a refractive index of 1.33 (or 1.465 for GdmCl) to the refractive index of 1.4 that we used.

| **Authors** | **Sample type** | **Dye** | **QY_D_** | κ^2^ | **Refractive index (RI)** | R_0_ **(Å)** | R_0_ **(Å) converted (for RI=1.4)** |
| --- | --- | --- | --- | --- | --- | --- | --- |
| **Cristóvão *et. al.***  **NAR**  **2012 (9)** | DNA | Alexa 488–C6-dT  Alexa 594–C6-dT | 0.6 | 2/3 | 1.33 (water) | 53.2 | 52.3 |
| **Reinartz *et.al.***  **J. Chem. Phys. 2018 (10)** | Protein | Alexa 488–C5-maleimide  Alexa 594–C5-maleimide | 0.92 | 2/3 | 1.33 (water) | 54 | 53 |
|  |  |  |  |  | 1.465 (GdmCl) | 51 | 52 |

**Table S3 – Gamma (detection factor) calculation:**

For each measurement day, the detection factor of the instrument was measured by using two FRET standards, 30 bp oligonucleotide with Alexa 488 (donor) and Alexa 594 (acceptor) either 10 bp (sample 10) or 21 bp (sample 21) apart. ALEX-FRET was performed and the proximity ratio P and stoichiometric value S were used as described in Lee et al. (2005) (2), to calculate the gamma factor. Using the experimentally derived gamma factor, and the observed proximity ratio, we calculated the inter-fluorophore distance in 10 bp and 21 bp samples (assuming R_o_ = 52 Å). As a test to check if the calculated gamma factor was correct, we used FRET Positioning and Screening (FPS) (11, 12), to calculate the Förster distance-independent inter-fluorophore distances (R_mp_ and <R_DA_>) between the AV simulated dyes. For 10 bp, R_mp_ = 44.5 Å, <R_DA_> = 47 Å. For 21 bp, R_mp_ = 67.5 Å, <R_DA_> = 69.5 Å (for definitions of R_mp_, <R_DA_>, see refs: 12 and 13). Our FRET-derived distances, R, between the dyes on the 10 bp and 21 bp samples fall between the theoretically calculated R_mp_ and <R_DA_> values.

| **Day of measurement** | **Proximity ratio Sample 10** | **Stoichiometry Sample 10** | **Proximity Ratio Sample 21** | **Stoichiometry Sample 21** | **γ** | **R sample 10 (Å)** | **R sample 21 (Å)** |
| --- | --- | --- | --- | --- | --- | --- | --- |
| 1 | 0.7 | 0.45 | 0.2 | 0.46 | 0.93 | 46 | 66 |
| 2 | 0.67 | 0.51 | 0.2 | 0.51 | 1.03 | 47 | 66 |
| 3 | 0.7 | 0.5 | 0.17 | 0.5 | 0.95 | 45 | 67 |
| 4 | 0.7 | 0.5 | 0.16 | 0.5 | 0.9 | 45 | 68 |
| 5 | 0.67 | 0.51 | 0.16 | 0.52 | 0.92 | 46 | 68 |
| 6 | 0.68 | 0.5 | 0.15 | 0.51 | 0.88 | 45 | 68 |
| 7 | 0.63 | 0.52 | 0.18 | 0.54 | 0.8 | 46 | 65 |
| 8 | 0.7 | 0.51 | 0.18 | 0.48 | 1.23 | 46 | 69 |

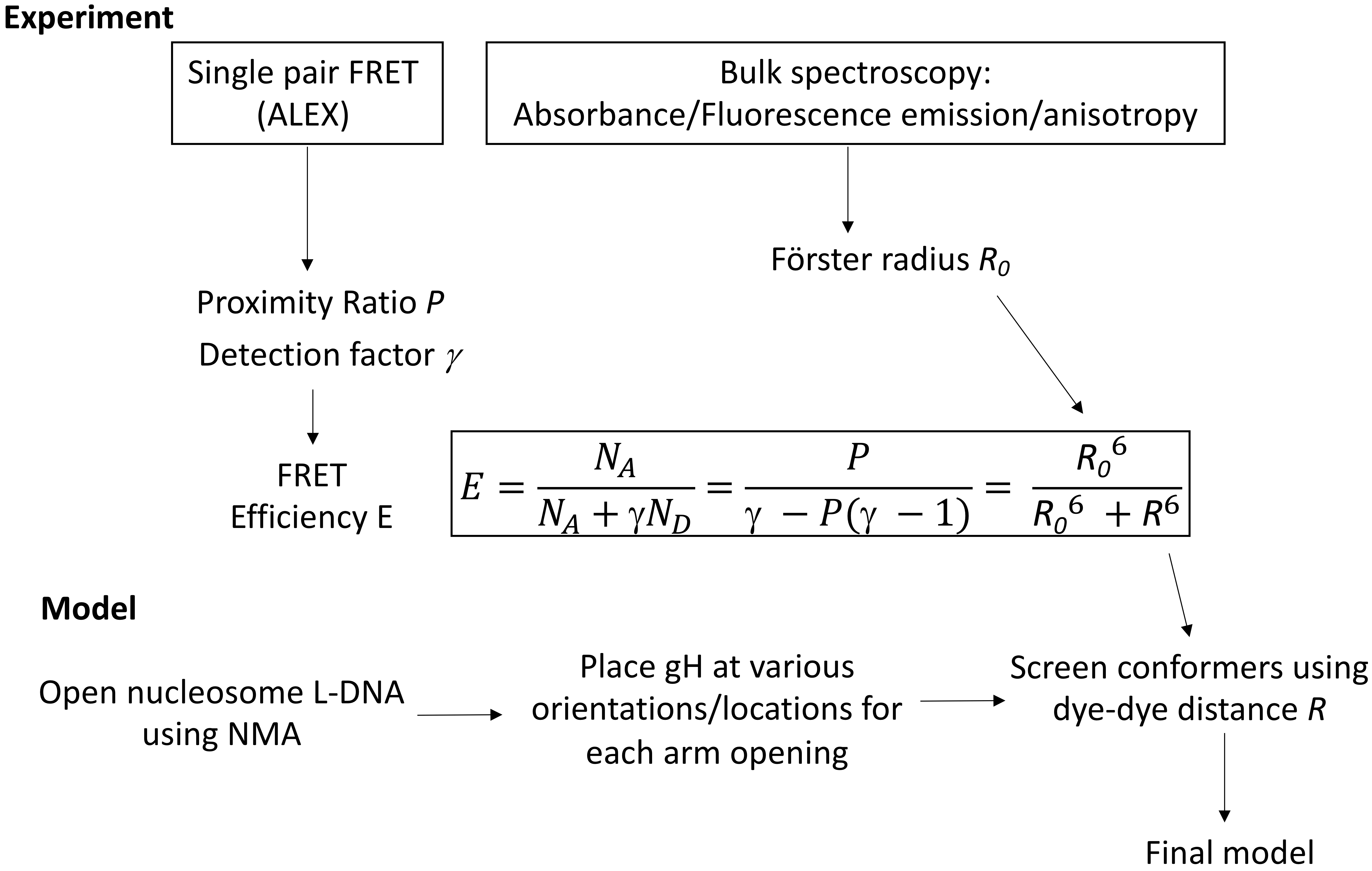

**Fig S5:** Workflow to derive the final structural models from the experimental data.

**Methods: Modelling nucleosomes based on experimental distances**

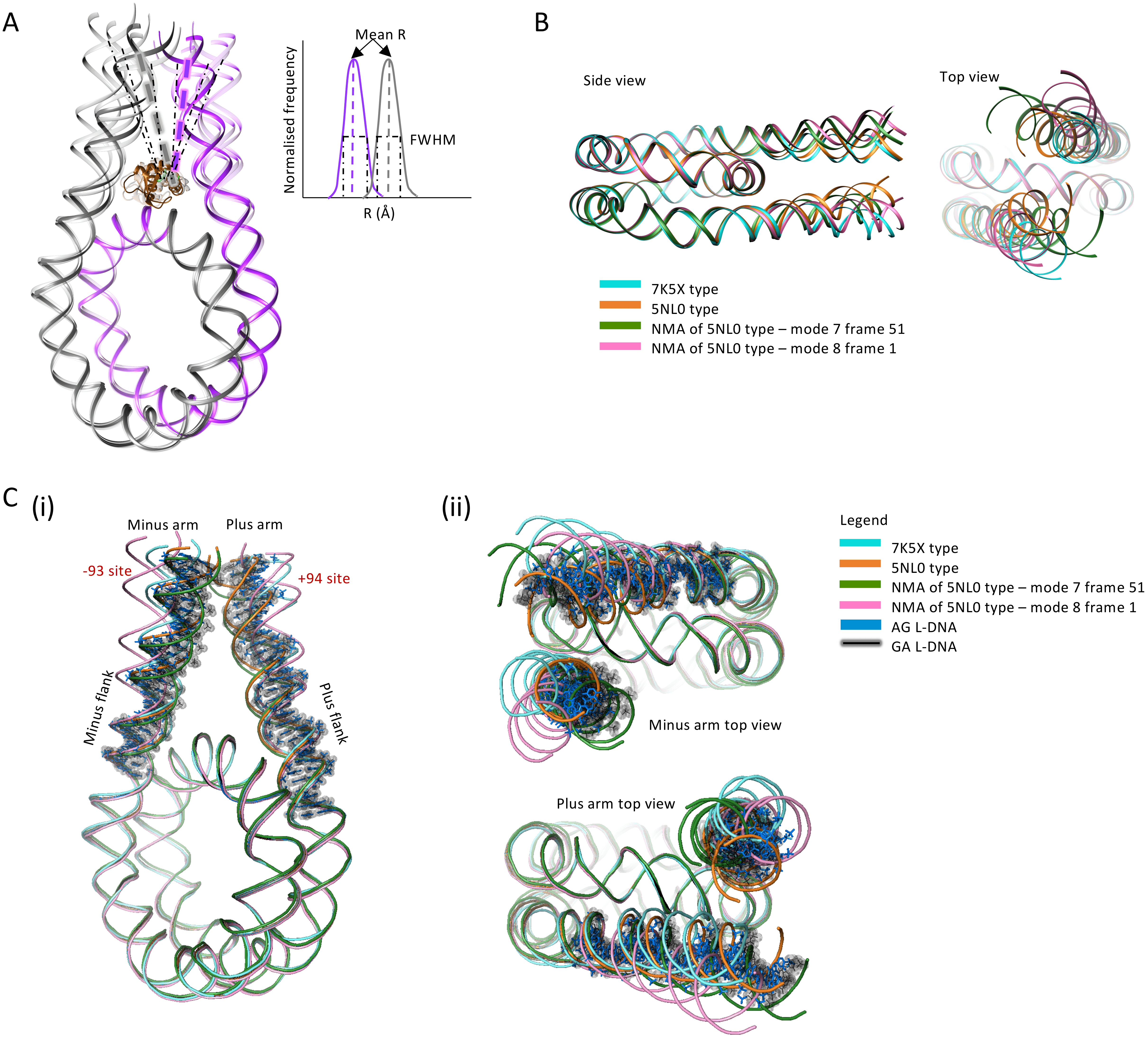

**Fig S6: A.** Schematic of the models of the gH-nucleosome complexes generated. The DNA is shown in backbone ribbon representation (gray: minus arm, purple: plus arm) and the gH is shown in brown. The core histones are not shown for clarity. The length of the DNA in the nucleosome models is 193bp instead of the 226bp used for experiments. The full-width at the half maxima (FWHM) of the FRET peaks can be explained by L-DNA arm movements and slight orientational changes in the gH. **B.** Four arm-opening types of structures of the nucleosomes were generated, the 7K5X-type (13) (cyan), the 5NL0-type (14) (orange), and two arm-opening structures obtained by subjecting the 5NL0-type structure to Elastic Network Normal Mode Analysis (15): mode 7 frame 51 (green) and mode 8 frame 1 (pink). **C. (i)** The four arm-opening types were aligned to AG (blue) and GA (black) L-DNA modeled using the cgDNA webserver (16). These show sequence-dependent bending. Since FRET distances were not computed for the cgDNA modeled L-DNAs, we align these models with the four arm-opening types studied. The alignment shows **(ii)** that the cgDNA modeled AG and GA arms fall within the range of arm-opening types studied.

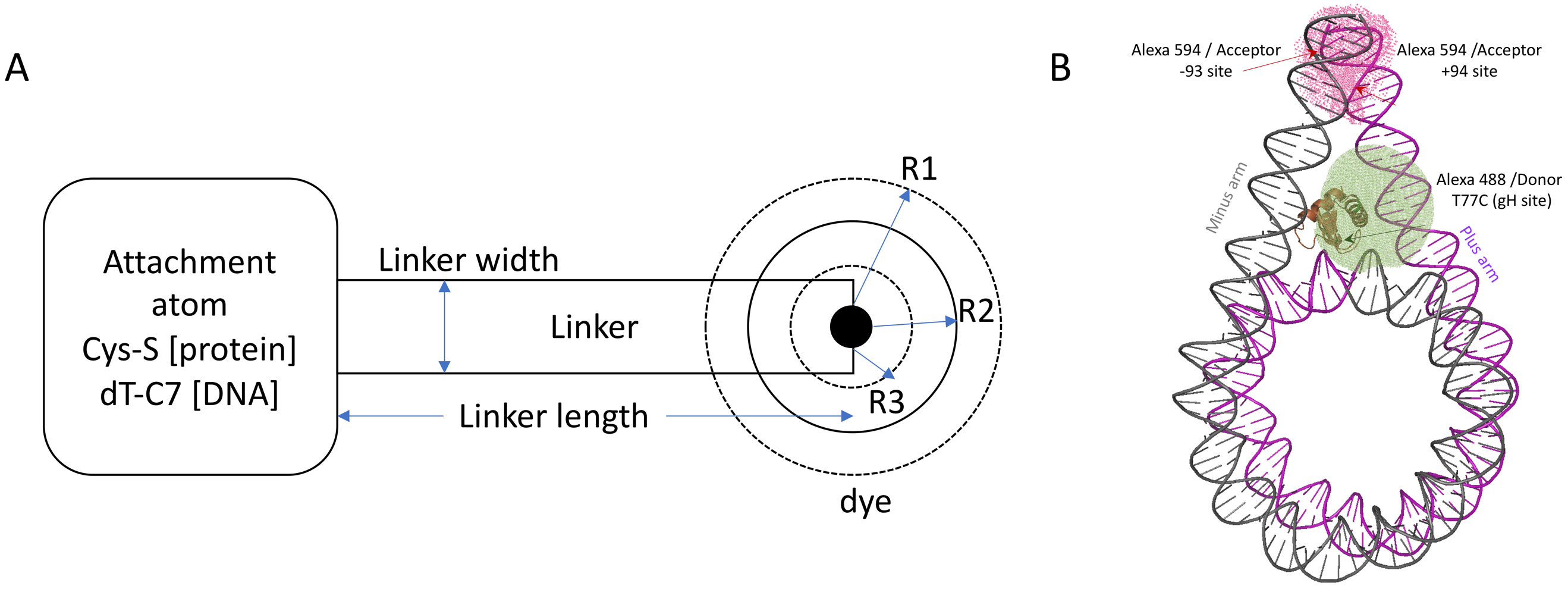

**Figure S7:** Accessible volume (AV) simulation of the fluorophores using FPS (12). **A.** Schematic of the dye adapted from Kalinin et al. (2012) (12) describing the parameters used for the AV simulations of the dyes (Table T7), such as the linker length and width, and the three dye radii, R1, R2 and R3. **B.** One nucleosome structure is shown [5NL0-type]: The purple arm is plus L-DNA and the grey arm is minus L-DNA. The accessible volume sphere for the Alexa 488 donor dye is shown in green and that of the Alexa 594 acceptor dye is shown in pink.

**Table S4 – Parameters used for accessible volume simulation of dyes:** We used the following dye parameters for screening structures with the FRET Positioning and Screening (FPS) software (12). The attachment points of the linkers to which the fluorophores are associated were the cysteine 77 sulphur atom on the gH for the donor dye, and the C7 atom of thymine +94 or -93 on the DNA for the acceptor dye.

| All values in Å | | | | | |
| --- | --- | --- | --- | --- | --- |
| **dye** | **Linker length** | **Linker width** | **R1** | **R2** | **R3** |
| Donor:  Alexa 488-C5-maleimide-Cys (S atom) [protein label]  (FPS 1.1 software (12)) | 20.5 | 4.5 | 5.0 | 4.5 | 1.5 |
| Acceptor:  Alexa 594-C6-dT (C7 atom)  [DNA label]  (17) | 20 | 4.5 | 8.1 | 3.2 | 2.6 |

**Results – Proximity Ratio plots:** All single-molecule experiments were performed with dual laser excitation (ALEX). In the following figures, each row A-D shows the raw 2D plot (Stoichiometry or S on y-axis, Proximity ratio P on x-axis) as obtained in ALEX experiments. The doubly labelled sample has a stoichiometry range from 0.25 to 0.75, peaking at 0.5. An S range of 0.25 to 0.75 and a P of 0 to 1 is selected (purple rectangle) and the proximity ratio histograms are fitted to multiple Gaussians (lower rows in A-D). On the right, all replicates and the averaged histogram (black trace) are plotted together.

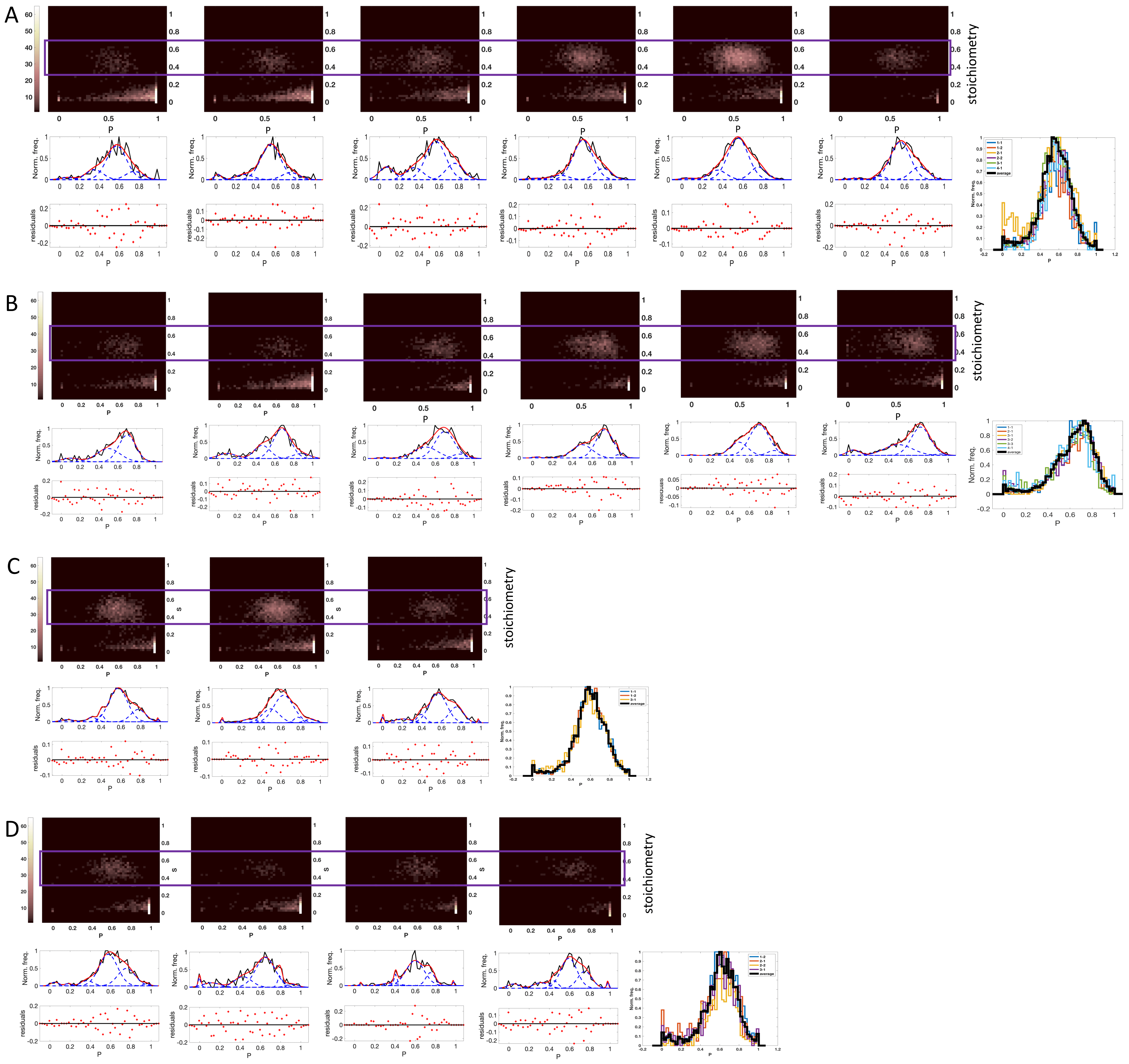

**Figure S8: A –** MG nucleosome labelled on the minus L-DNA arm at -93 with Alexa 594, and on the LH at T77C (gH site) with Alexa 488. **B –** MG nucleosome labelled on plus L-DNA arm at +94 with Alexa 594, and on the LH at T77C (gH site) with Alexa 488. **C –** GM nucleosome labelled on minus L-DNA arm at -93 with Alexa 594, and on the LH at T77C (gH site) with Alexa 488. **D –** GM nucleosome labelled on plus L-DNA arm at +94 with Alexa 594, and on the LH at T77C (gH site) with Alexa 488.

**Table S5 Proximity ratio from replicates of constructs MG and GM labelled at L-DNA arms and the gH domain of the LH:** Mean proximity ratio (obtained by fitting the P histograms to multiple Gaussians) of the major population peak are shown here. Standard deviations are calculated from the mean proximity ratio values. The full width at half maximum and the relative area of the peaks are given. Data shown are from replicates and from the plot averaged from the replicates.

| Constructs | MG (-) T77C | MG (+) T77C | GM (-) T77C | GM (+) T77C |
| --- | --- | --- | --- | --- |
| Mean P of peak(s) of replicates | 0.57 | 0.70 |  |  |
|  | 0.55 | 0.67 | 0.57 | 0.58 |
|  | 0.57 | 0.70 | 0.60 | 0.65 |
|  | 0.55 | 0.72 | 0.56 | 0.60 |
|  | 0.55 | 0.70 |  | 0.60 |
|  | 0.56 | 0.72 |  |  |
| Standard deviation of mean P from replicates | ±0.01 | ±0.02 | ±0.02 | ±0.03 |
| Mean P of peak in averaged plot | 0.55 | 0.72 | 0.57 | 0.60 |
| Full-width at half maximum of peak(s) of replicates | 0.33 | 0.30 |  |  |
|  | 0.33 | 0.30 | 0.30 | 0.30 |
|  | 0.33 | 0.33 | 0.33 | 0.32 |
|  | 0.33 | 0.29 | 0.30 | 0.30 |
|  | 0.33 | 0.33 |  | 0.26 |
|  | 0.33 | 0.32 |  |  |
| Full-width at half maximum of peak in averaged plot | 0.33 | 0.33 | 0.32 | 0.28 |
| Relative area under the peak of replicates (%) | 66.0 | 55.0 |  |  |
|  | 74.0 | 58.0 | 68.0 | 58.0 |
|  | 53.0 | 60.0 | 60.0 | 61.0 |
|  | 73.0 | 63.0 | 58.0 | 68.0 |
|  | 69.0 | 67.2 |  | 52.3 |
|  | 73.0 | 58.5 |  |  |
| Relative area under the peak in averaged plot (%) | 67.0 | 63.0 | 67.0 | 57.0 |

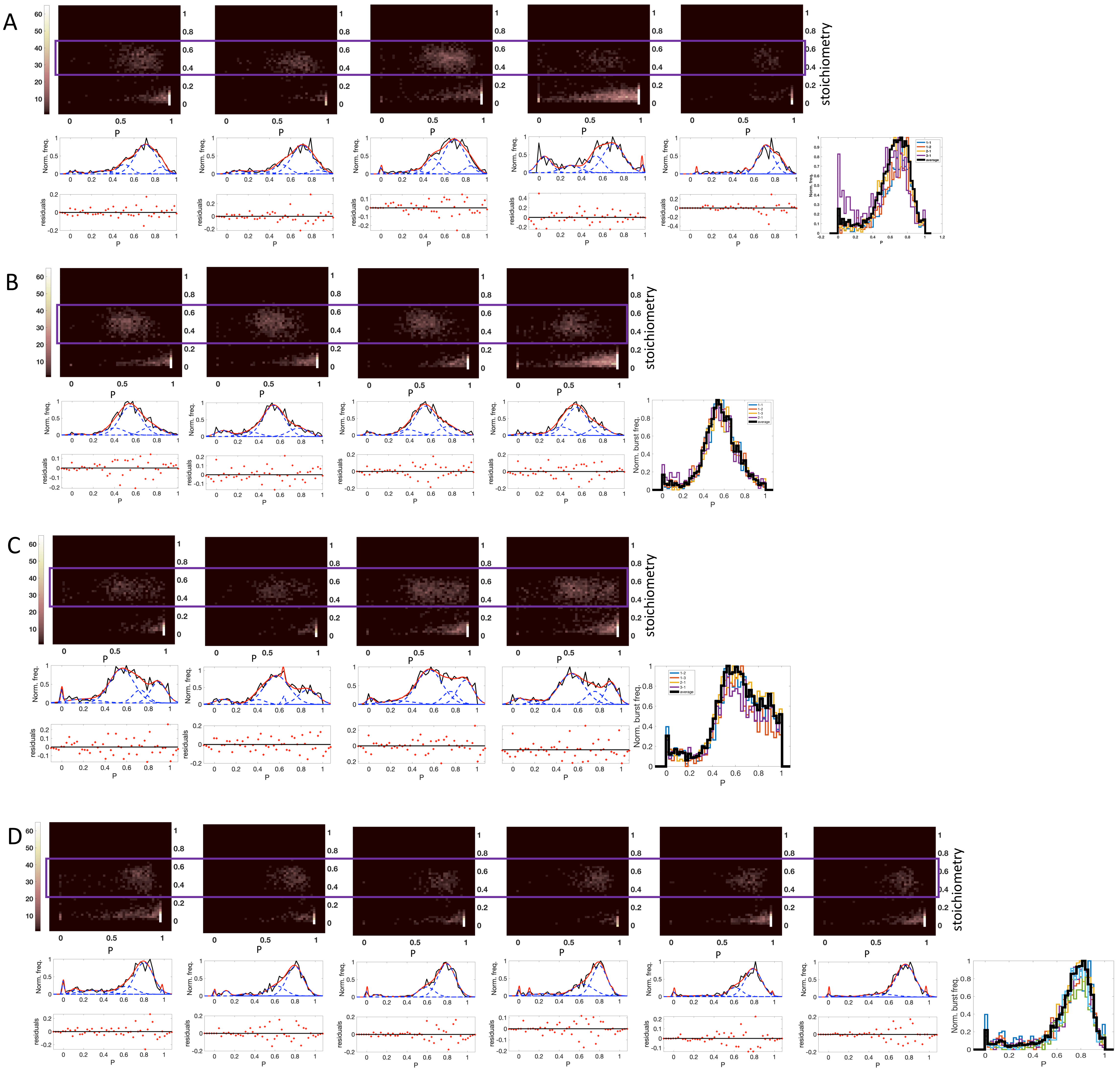

**Figure S9: A –** AG nucleosome labelled on minus L-DNA arm at -93 with Alexa 594, and on the LH at T77C (gH site) with Alexa 488. **B –** AG nucleosome labelled on plus L-DNA arm at +94 with Alexa 594, and on the LH at T77C (gH site) with Alexa 488. **C –** GA nucleosome labelled on minus L-DNA arm at -93 with Alexa 594, and on the LH at T77C (gH site) with Alexa 488. **D –** GA nucleosome labelled on plus L-DNA arm at +94 with Alexa 594, and on the LH at T77C (gH site) with Alexa 488.

**Table S6 Proximity ratio from replicates of constructs AG and GA labelled at L-DNA arms and the gH domain of the LH:** Mean proximity ratio (obtained by fitting the P histograms to multiple Gaussians) of the major population peak are shown here. In case a second peak is reproducibly present, it is shown as well. Standard deviations are calculated from the mean proximity ratio values. The full-width at half maximum and the relative area of the peaks are given. Data shown are from replicates and from the plot averaged from the replicates.

| Constructs | AG (-) T77C | AG (+) T77C | GA (-) T77C | | GA (+) T77C |
| --- | --- | --- | --- | --- | --- |
| Mean P of peak(s) of replicates | 0.7 |  | major | minor | 0.80 |
|  | 0.71 | 0.55 | 0.55 | 0.89 | 0.80 |
|  | 0.71 | 0.55 | 0.57 | 0.86 | 0.79 |
|  | 0.7 | 0.55 | 0.56 | 0.89 | 0.80 |
|  | 0.71 | 0.54 | 0.54 | 0.9 | 0.79 |
|  |  |  |  | | 0.77 |
| Standard deviation of mean P from replicates | ±0.005 | ±0.005 | ±0.01 | ±0.02 | ±0.01 |
| Mean P of peak in averaged plot | 0.71 | 0.54 | 0.55 | | 0.80 |
| Full-width at half maximum of peak(s) of replicates | 0.33 |  | major | minor | 0.30 |
|  | 0.33 | 0.33 | 0.37 | 0.24 | 0.27 |
|  | 0.33 | 0.33 | 0.39 | 0.29 | 0.28 |
|  | 0.33 | 0.33 | 0.39 | 0.24 | 0.25 |
|  | 0.24 | 0.33 | 0.39 | 0.19 | 0.27 |
|  |  |  |  | | 0.32 |
| Full-width at half maximum of peak in averaged plot | 0.33 | 0.33 | 0.39 | 0.22 | 0.27 |
| Relative area under the peak of replicates (%) | 69.0 |  | major | minor | 64.0 |
|  | 67.0 | 65.0 | 55.0 | 22.0 | 66.0 |
|  | 56.0 | 61.0 | 61.0 | 23.0 | 74.0 |
|  | 51.0 | 67.0 | 54.0 | 24.0 | 57.6 |
|  | 68.0 | 66.0 | 57.0 | 19.0 | 71.0 |
|  |  |  |  | | 83.0 |
| Relative area under the peak in averaged plot (%) | 58.0 | 66.0 | 57.0 | 21.0 | 62.0 |

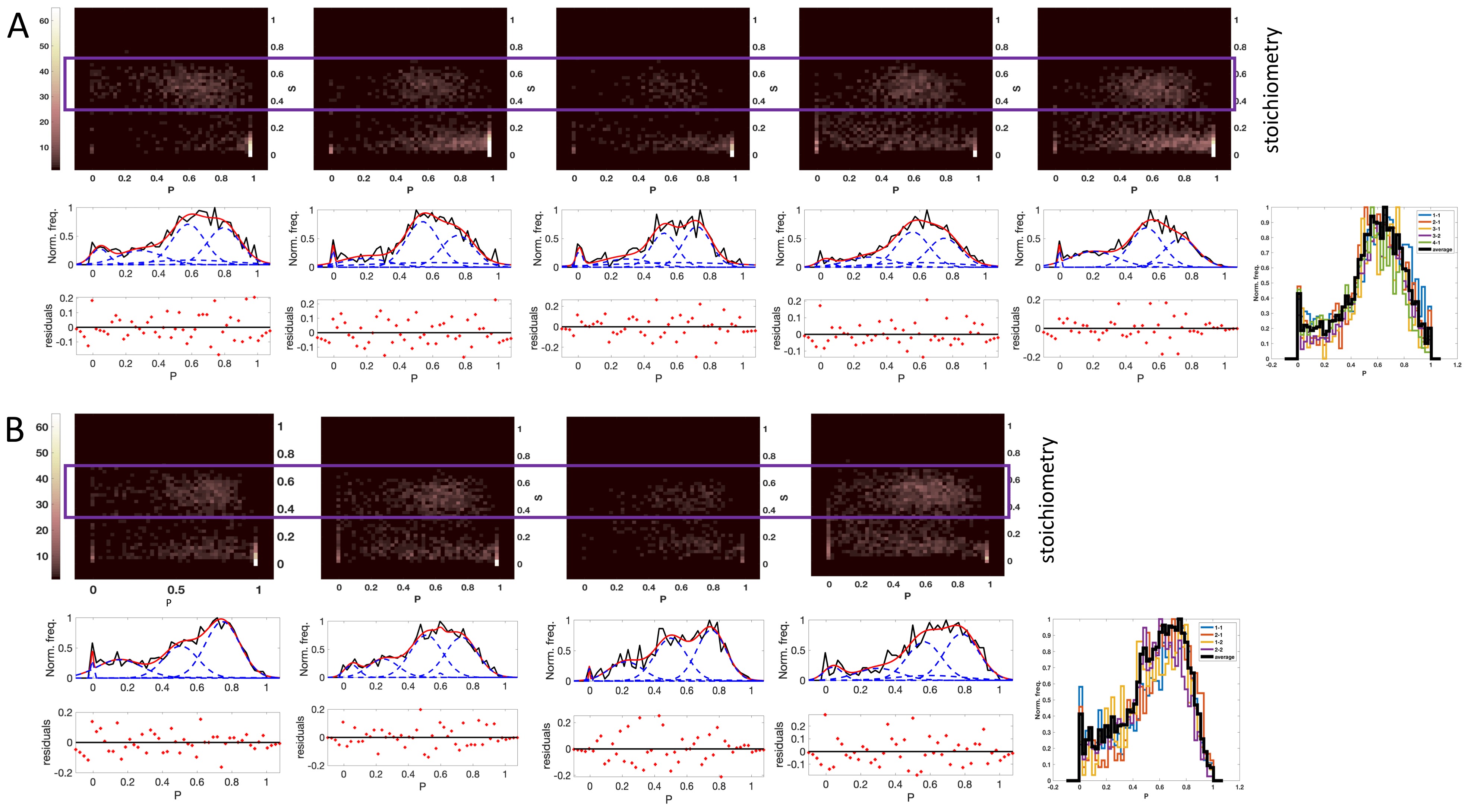

**Figure S10: A –** TU nucleosome labelled on minus L-DNA arm at -93 with Alexa 594, and on the LH at T77C (gH site) with Alexa 488. **B –** TU nucleosome labelled on plus L-DNA arm at +94 with Alexa 594, and on the LH at T77C (gH site) with Alexa 488.

**Table S7 Proximity ratio from replicates of constructs TU labelled at L-DNA arms and the gH domain of the LH:** Mean proximity ratio (obtained by fitting the P histograms to multiple Gaussians) of the two peaks are shown here. Standard deviations are calculated from the mean proximity ratio values. The full-width at half maximum and the relative area of the peaks are given. Data shown are from replicates and from the plot averaged from the replicates.

| Constructs | TU (-) T77C | | TU (+) T77C | |
| --- | --- | --- | --- | --- |
| Mean P of peak(s) of replicates |  |  |  |  |
|  | 0.58 | 0.79 | 0.51 | 0.75 |
|  | 0.54 | 0.75 | 0.57 | 0.78 |
|  | 0.52 | 0.72 | 0.50 | 0.75 |
|  | 0.56 | 0.75 | 0.51 | 0.73 |
|  | 0.52 | 0.73 |  |  |
| Standard deviation of mean P from replicates | ±0.03 | ±0.03 | ±0.03 | ±0.02 |
| Mean P of peak in averaged plot | 0.54 | 0.75 | 0.52 | 0.74 |
| Full-width at half maximum of peak(s) of replicates |  |  |  |  |
|  | 0.33 | 0.33 | 0.33 | 0.33 |
|  | 0.33 | 0.33 | 0.33 | 0.33 |
|  | 0.26 | 0.28 | 0.33 | 0.30 |
|  | 0.33 | 0.33 | 0.33 | 0.33 |
|  | 0.33 | 0.33 |  |  |
| Full-width at half maximum of peak in averaged plot | 0.33 | 0.33 | 0.33 | 0.33 |
| Relative area under the peak of replicates (%) | 35.0 | 32.0 | 30.0 | 50.0 |
|  | 42.0 | 30.0 | 33.0 | 40.0 |
|  | 28.0 | 36.0 | 38.0 | 41.0 |
|  | 37.0 | 31.0 | 40.0 | 38.0 |
|  | 41.5 | 30.4 |  |  |
| Relative area under the peak in averaged plot (%) | 38 | 31 | 31 | 40 |

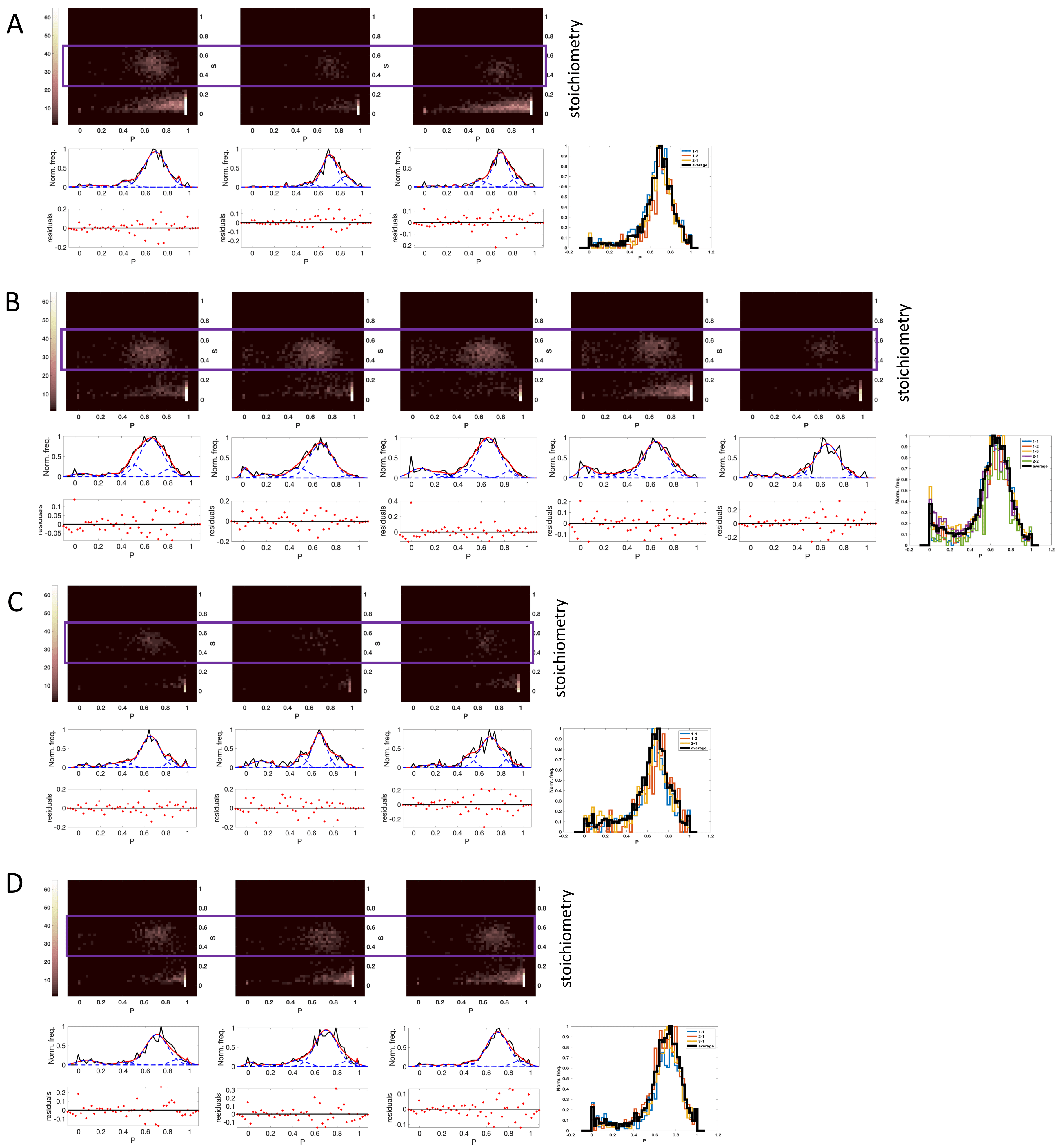

**Figure S11: A –** MG nucleosome labelled on minus L-DNA arm at -93 with Alexa 594, and on the LH at G101C (CTD site) with Alexa 488. **B –** MG nucleosome labelled on plus L-DNA arm at +94 with Alexa 594, and on the LH at G101C (CTD site) with Alexa 488. **C –** GM nucleosome labelled on minus L-DNA arm at -93 with Alexa 594, and on the LH at G101C (CTD site) with Alexa 488. **D –** GM nucleosome labelled on plus L-DNA arm at +94 with Alexa 594, and on the LH at G101C (CTD site) with Alexa 488.

**Table S8 Proximity ratio from replicates of constructs MG and GM labelled at L-DNA arms and the CTD domain of the LH:** Mean proximity ratio (obtained by fitting the P histograms to multiple Gaussians) of the major population peak are shown here. Standard deviations are calculated from the mean proximity ratio values. The full-width at half maximum and the relative area of the peaks are given. Data shown are from replicates and from the plot averaged from the replicates.

| Constructs | MG (-) G101C | MG (+) G101C | GM (-) G101C | GM (+) G101C |
| --- | --- | --- | --- | --- |
| Mean P of peak(s) of replicates |  | 0.66 |  |  |
|  | 0.69 | 0.67 | 0.66 | 0.69 |
|  | 0.70 | 0.66 | 0.70 | 0.70 |
|  | 0.69 | 0.64 | 0.67 | 0.68 |
|  |  | 0.66 |  |  |
| Standard deviation of mean P from replicates | ±0.006 | ±0.01 | ±0.02 | ±0.01 |
| Mean P of peak in averaged plot | 0.69 | 0.67 | 0.68 | 0.70 |
| Full-width at half maximum of peak(s) of replicates |  | 0.33 |  |  |
|  | 0.33 | 0.33 | 0.28 | 0.33 |
|  | 0.22 | 0.33 | 0.30 | 0.33 |
|  | 0.24 | 0.33 | 0.21 | 0.33 |
|  |  | 0.33 |  |  |
| Full-width at half maximum of peak in averaged plot | 0.27 | 0.33 | 0.26 | 0.33 |
| Relative area under the peak of replicates (%) |  | 70.0 |  |  |
|  | 84.0 | 68.0 | 71.3 | 74.0 |
|  | 69.0 | 64.0 | 69.0 | 74.0 |
|  | 68.0 | 67.0 | 54.3 | 79.0 |
|  |  | 83.0 |  |  |
| Relative area under the peak in averaged plot (%) | 73.0 | 66.0 | 60.0 | 77.0 |

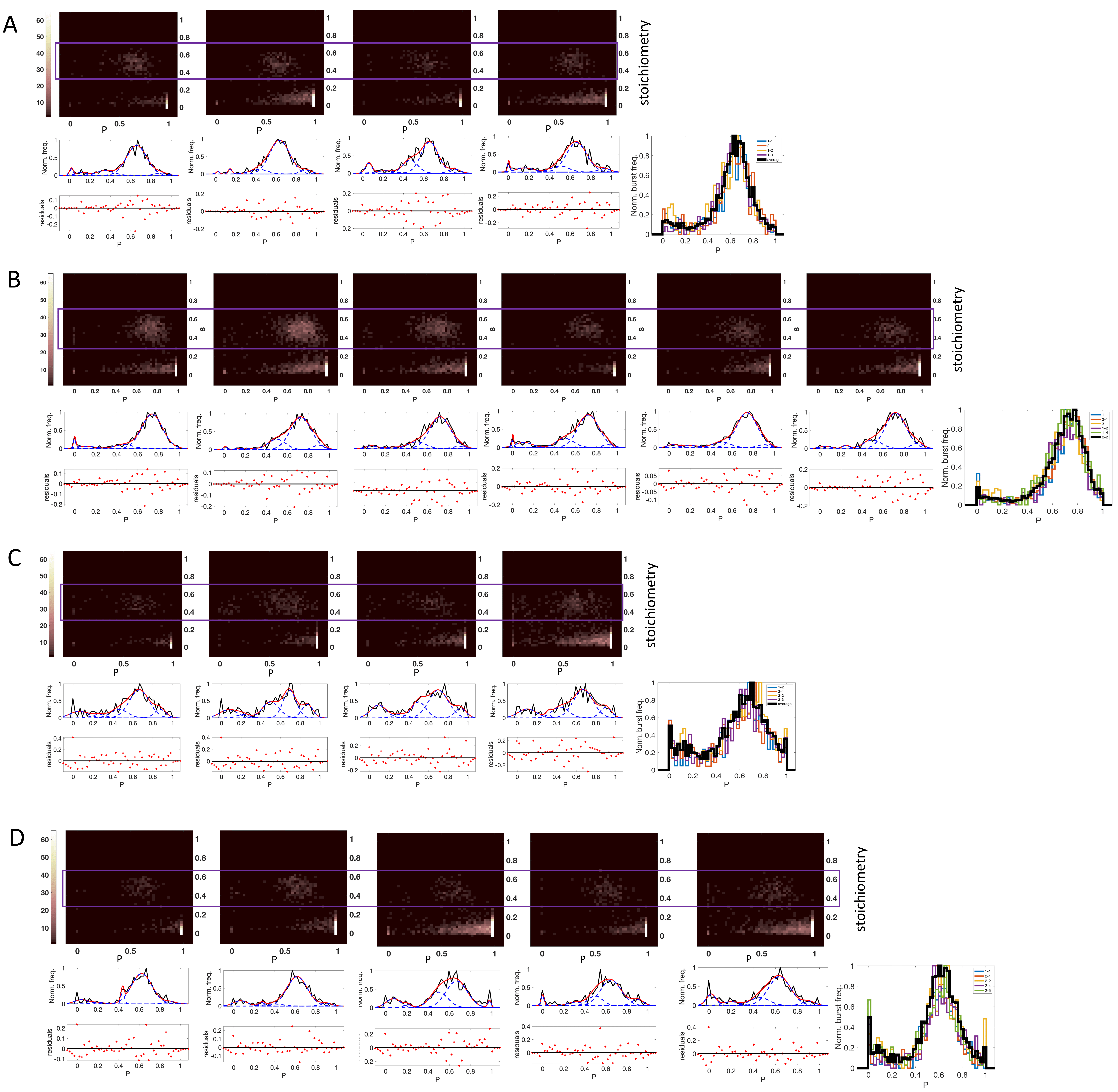

**Figure S12: A –** AG nucleosome labelled on minus L-DNA arm at -93 with Alexa 594, and on the LH at G101C (CTD site) with Alexa 488. **B –** AG nucleosome labelled on plus L-DNA arm at +94 with Alexa 594, and on the LH at G101C (CTD site) with Alexa 488. **C –** GA nucleosome labelled on minus L-DNA arm at -93 with Alexa 594, and on the LH at G101C (CTD site) with Alexa 488. **D –** GA nucleosome labelled on plus L-DNA arm at +94 with Alexa 594, and on the LH at G101C (CTD site) with Alexa 488.

**Table S9 Proximity ratio from replicates of constructs AG and GA labelled at L-DNA arms and the CTD domain of the LH:** Mean proximity ratio (obtained by fitting the P histograms to multiple Gaussians) of the major population peak are shown here. In case a second peak is present, it is shown as well. Standard deviations are calculated from the mean proximity ratio values. The full-width at half maximum and the relative area of the peaks are given. Data shown are from replicates and from the plot averaged from the replicates.

|  | AG (-) G101C | AG (+) G101C | GA (-) G101C | | GA (+) G101C |
| --- | --- | --- | --- | --- | --- |
| Mean P of peak(s) of replicates |  | 0.74 |  | | 0.62 |
|  | 0.66 | 0.74 | 0.66 | | 0.62 |
|  | 0.66 | 0.72 | 0.50 | 0.70 | 0.67 |
|  | 0.66 | 0.72 | 0.70 | | 0.65 |
|  | 0.64 | 0.73 | 0.67 | | 0.64 |
|  |  | 0.72 |  | |  |
| Standard deviation of mean P from replicates | ±0.01 | ±0.01 | ±0.02 | | ±0.02 |
| Mean P of peak in averaged plot | 0.65 | 0.73 | 0.70 | | 0.65 |
| Full-width at half maximum of peak(s) of replicates |  | 0.33 |  | | 0.33 |
|  | 0.33 | 0.33 | 0.33 | | 0.33 |
|  | 0.33 | 0.33 | 0.28 | 0.22 | 0.33 |
|  | 0.28 | 0.33 | 0.33 | | 0.33 |
|  | 0.33 | 0.33 | 0.33 | | 0.33 |
|  |  | 0.33 |  | |  |
| Full-width at half maximum of peak in averaged plot | 0.33 | 0.33 | 0.33 | | 0.33 |
| Relative area under the peak of replicates (%) |  | 84.0 |  | | 81.0 |
|  | 78.0 | 78.0 | 57.6 | | 78.0 |
|  | 71.0 | 70.0 | 28.0 | 37.0 | 55.0 |
|  | 60.0 | 77.0 | 48.0 | | 64.0 |
|  | 81.0 | 71.0 | 54.0 | | 63.0 |
|  |  | 74.0 |  | |  |
| Relative area under the peak in averaged plot (%) | 76.0 | 76.0 | 49.5 | | 61.0 |

**Results – Distance plots**

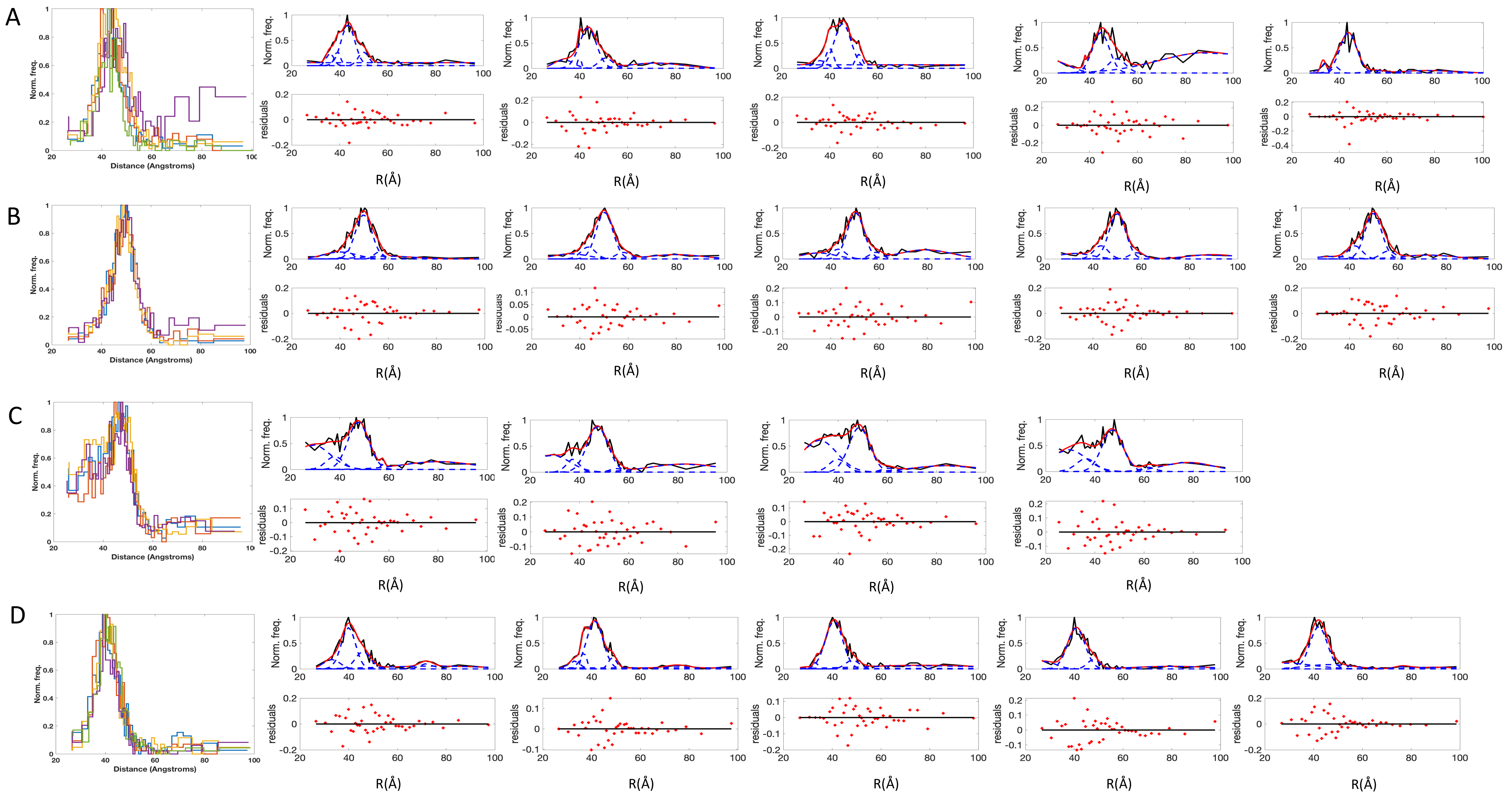

**Figure S13:** Distance histograms fitted to multiple Gaussians. Data are shown for 4-5 replicas for four nucleosome constructs. **A:** AG nucleosome labelled with the donor dye at the LH gH domain (T77C site), and the acceptor dye at -93 site on the minus L-DNA arm. **B:** AG nucleosome labelled with the donor dye at the LH gH domain (T77C site), and the acceptor dye at +94 site on the plus L-DNA arm. **C:** GA nucleosome labelled with the donor dye at the LH gH domain (T77C site), and the acceptor dye at -93 site on the minus L-DNA arm. **D:** GA nucleosome labelled with the donor dye at the LH gH domain (T77C site), and the acceptor dye at +94 site on the plus L-DNA arm.

**Table S10 – Interfluorophore distances, R (in Å) from experiments:** Mean distance of the major peak in distance histograms (Fig. S6). Data are shown for replicates. The detection factor or γ value for that measurement day is given in [] brackets. The average and standard deviation are obtained from replicates.

| Constructs: | AG (-) T77C | AG (+) T77C | GA (-) T77C | GA (+) T771C |
| --- | --- | --- | --- | --- |
| Mean R of peak(s) of replicates (Å)  and [γ value] | 43.5 [0.95] |  |  | 40.0 [0.88] |
|  | 43.5 [0.95] | 49.7 [0.90] | 49.0 [0.92] | 41.3 [0.88] |
|  | 45.4 [0.96] | 50.0 [0.90] | 48.6 [0.92] | 40.7 [0.92] |
|  | 44.7 [1.03] | 50.0 [0.90] | 50.0 [0.95] | 40.8 [0.91] |
|  | 43.8 [1.23] | 50.3 [0.93] | 48.3 [0.8] | 42.0 [0.95] |
| Standard deviation of mean R (Å) from replicates | ±0.8 | ±0.2 | ±0.74 | ±0.7 |
| Average of Mean R (Å) from replicates | 44.2 | 50.0 | 49.0 | 41.0 |
| Full-width at half maximum (Å) of peak(s) of replicates | 9.6 |  |  | 10.0 |
|  | 12.0 | 11.5 | 14.0 | 11.0 |
|  | 12.0 | 11.5 | 14.0 | 12.0 |
|  | 12.0 | 11.5 | 14.0 | 12.0 |
|  | 12.0 | 11.5 | 14.0 | 12.0 |

**Table S11: Results – Computed distances from models:** Distances obtained from models with Zone B close to the A-tract. The corresponding models are shown in Figure 3 in the main manuscript. The experiment-derived distances are added for comparison. The experimental half-width at half maximum (HWHM) of the major peaks are given as a measure of deviation from the mean. For the AG construct, computed distances from three configurations per arm-opening are shown, with their averages and standard deviations. For the GA construct, the computed distances from two configurations per arm-opening are shown but not averaged. Numbers in bold are the model distances that are closest to the experimental distances. The corresponding models are shown in Figure 3 in main text.

| All values are in Å | | | | |
| --- | --- | --- | --- | --- |
| Constructs | AG (-) T77C | AG (+) T77C | GA (-) T77C | GA (+) T77C |
| Experiment: Avg. of mean R from replicates ± SD of mean R from replicates | 44.2±0.8 | 50.0±0.2 | 49.0±0.74 | 41.0±0.7 |
| Experiment: half-width at half maxima | 6.0 | 6.0 | 7.0 | 6.0 |
| Model: 5NL0-type | 46.0±1.6 | 53.0±1.6 | 47.4 | 42.5 |
|  | **44.6** | **51.1** |  |  |
|  | 46.6 | 53.3 | 54.4 | 48.5 |
|  | 47.7 | 54.3 |  |  |
| Model: 7K5X-type | 50.0±1.6 | 56.0±1.5 | 44.9 | 42.8 |
|  | 48.2 | 54.4 |  |  |
|  | 50.3 | 57.3 | 53.0 | 50.0 |
|  | 51.6 | 56.6 |  |  |
| Model: mode 7 frame 51 | 46.3±1.6 | 56.0±1.6 | **50.2** | **40.6** |
|  | 43.5 | 54.0 |  |  |
|  | 45.0 | 56.2 | 58.0 | 46.0 |
|  | 46.3 | 57.0 |  |  |
| Model: mode 8 frame 1 | 50.2±1.2 | 55.4±1.6 | 50.0 | 46.0 |
|  | 50.2 | 53.6 |  |  |
|  | 52.0 | 56.5 |  |  |
|  | 54.1 | 56.0 |  |  |

**Alternate models of nucleosomes I: R74 proximal to A-tract**

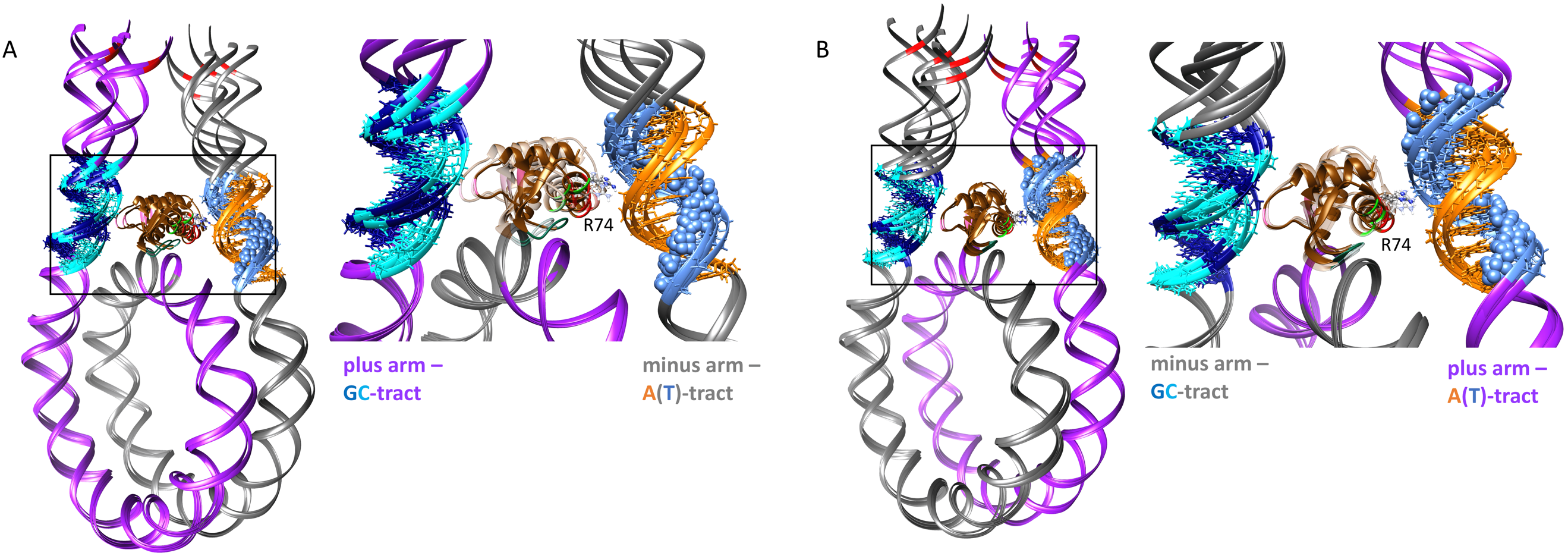

**Figure S14: A.** AG construct and **B.** GA construct. In both, the Zone B α3 helix is positioned close to the A-tract minor groove. Insets show R74 close to the minor groove of the A-tracts (in blue and orange). The structural ensembles were built using four arm openings, as described in Supplementary Material section S5. For each arm-opening, the gH was placed with the zone A (R74) proximal to the A-tract minor groove.

**Table S12**: **Comparison between distances from experiments and from the alternate models with R74 proximal to the A-tract** . The experimental mean R values (in Å) of the major peaks are given along with the standard deviation of the mean R values between replicates. For the models, computed distances from the gH orientations are averaged for each arm-opening and their standard deviations are calculated. Distances from individual models are not given. The experimental half-widths at half maximum (HWHM) of the major peaks are given as a measure of the deviation from the mean. The experimental distance peaks have a half-width at half maximum of ± 6 - 7Å. However, the R values for the A-tract arms (AG minus and GA plus) in the alternate models differ by 15 - 20 Å from the experimental mean R values.

| All values are in Å | | | | |
| --- | --- | --- | --- | --- |
| Constructs | AG (-) T77C | AG (+) T77C | GA (-) T77C | GA (+) T77C |
| Experiment: mean R of peak of averaged data ± SD of mean R from replicates | 44.2±0.8 | 50.0±0.2 | 49.0±0.74 | 41.0±0.7 |
| Experiment: half-width at half maxima | 6.0 | 6.0 | 7.0 | 6.0 |
| Model: 5NL0-type | 60.0±1.0 | 54.0±1.0 | 52.0±1.6 | 59.0±1.4 |
| Model: 7K5X-type | 59.0±1.0 | 56.0±0.6 | 56.0±1.7 | 61.0±1.4 |
| Model: mode 7 frame 51 | 63.0±1.1 | 51.0±1.0 | 49.2±1.5 | 61.4±1.3 |
| Model: mode 8 frame 1 | 62.5±1.6 | 59.4±1.0 | 59.0±1.8 | 61.0±1.6 |

**Alternate models of nucleosomes II: Off-dyad positioning of gH**

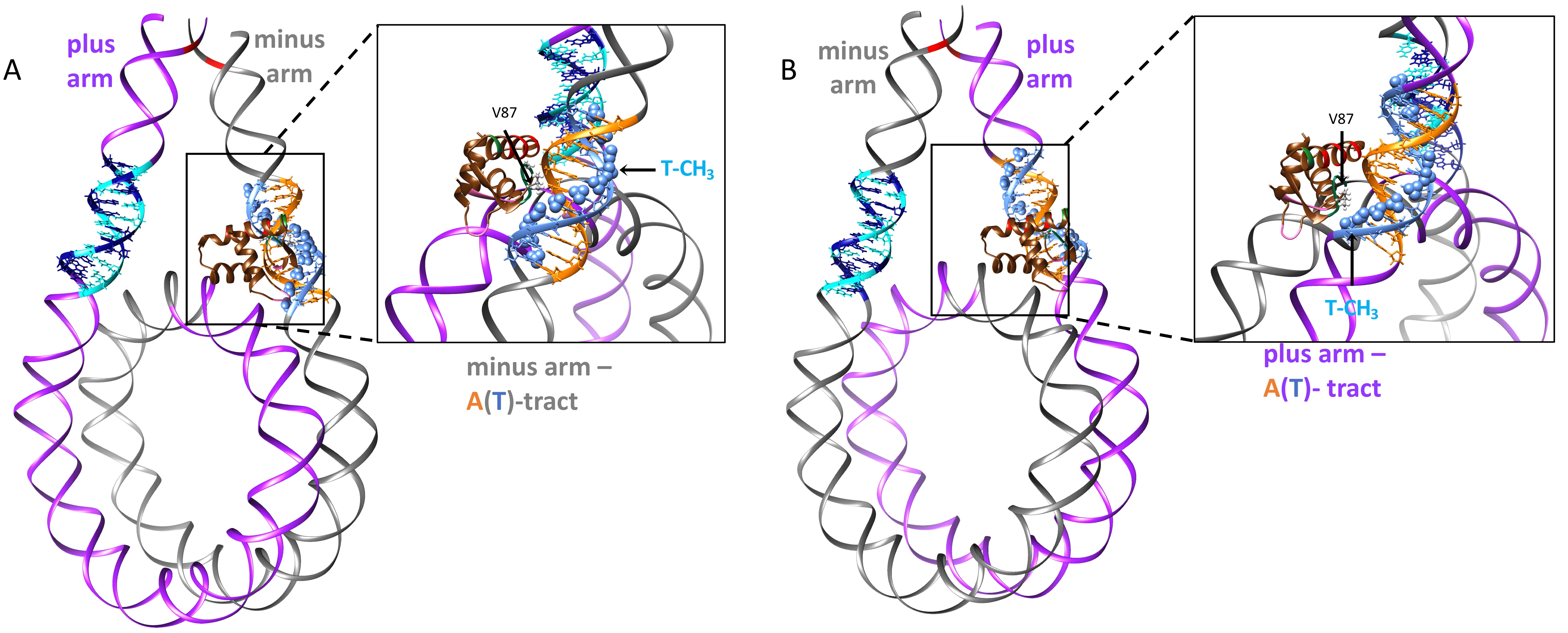

**Figure S15: A.** AG construct with the gH positioned off-dyad and proximal to the major groove of the A-tract on the grey minus arm. **B.** GA construct with the gH positioned off-dyad and proximal to the major groove of the A-tract on the purple plus arm. The boxed areas are shown in the insets to the right of each image, showing that V87 of the gH GVGA motif is proximal to the thymine methyl groups (blue spheres).

**Table S13:** **Comparison between distances from experiments and from the alternate models the gH placed off-dyad.** The distances between the fluorophore on the gH (T77C) and the A-tract containing arm (minus in AG, plus in GA) from experiment are compared to the distances obtained from the off-dyad models. The half-width at half-maximum of the major peaks shows a distance difference of about ± 6Å from the mean R value of the major peaks. However, the R values for the A-tract arms and the off-dyad positioned gH differ by +12 -16Å from the experimental mean R values.

| All values are in Å | | |
| --- | --- | --- |
| Constructs | AG (-) T77C | GA (+) T77C |
| Experiment: mean R of peak of averaged data ± SD of mean R from replicates | 44.2±0.8 | 41.0±0.7 |
| Experiment: half-width at half maxima | 6.0 | 6.0 |
| Model: 5NL0-type | 56.0 | 57.0 |

**Table S14: DNA proximal residues on the gH:** Comparison between X-ray crystal structures: 5NL0 (14), 4QLC(18), 5WCU(19), and FRET-restrained models: AG & GA.

|  | DNA-proximal residues in canonical gH binding mode (on-dyad) [from PDB: 5NL0(15), 4QLC(18), 5WCU(19)] | DNA-proximal residues in AG FRET-restrained models | DNA-proximal residues in GA FRET-restrained models |
| --- | --- | --- | --- |
| **Zone A** | K27 (T27 in 4QLC/5WCU), S29, Q67, R74 | E62, N63, Q67, R74 | E62, N63, S66, Q67 |
| **Zone B** | R42, R94 | R42, R94 (minor groove of A-tract linker DNA) | R42, R94 (minor groove of A-tract linker DNA), K40, K82 (backbone) |
| **Zone C** | S46, R47, Q48, S49, Q51, K52, K69, K73 (R73 in 4QLC/5WCU), K85, V87, A89, S90, S92 | S46, R47, Q48, S49, K52, K69, K85, V87, A89, S90, S92 | E39, S41, S43, S45, S46, R47, Q48, S49, Q51, K52, Y53, K55, N56, H57, D65, K69, K85, V87, A89, S90, S92 |

**Table S15: Sequences of Zones A, B, and C from LH isoforms from different organisms**:

Sequences of all LH isoforms shown were obtained from the Uniprot database (https://www.uniprot.org/). All the isoforms were modelled using Swiss-Model (swissmodel.expasy.org) and the Zones A, B, and C sequences were extracted. The arginines that we suggest to be responsible for the A-tract recognition, present in loop 1 and β2 of zone B, are depicted in pink bold letters.

The GVGA motif in Zone C, or the hydrophobic patch, is indicated in bold italics.

The α3 helix (zone A) is shown in its entirety. The arginine suggested to interact with the minor grooves of AT-rich regions (13) is denoted in bold red. Other arginines, present elsewhere on the alpha helix, are denoted in bold. Only 2 isoforms are shown for *M. musculus* and only one for *G. gallus* and *D. melanogaster*, respectively.

| Organism | LH isoforms | helix α3  [**Zone A**] | Loop 1  [**Zone B**] | β2  [**Zone B**] | beta-hairpin loop  [**Zone C**] |
| --- | --- | --- | --- | --- | --- |
| *Xenopus laevis* | H1A | VDKNNS**R**LKLALK | KE**R**SG | SFKL | GSGA |
|  | H1B | NS**R**LKLALKALVTK | KE**R**SG | SFKL | GSGA |
|  | H1C | VDKNNS**R**LKLALK | KE**R**GG | SFKL | GSGA |
|  | H1.0-A | DSQIKLSIK**R**LV | KS**R**SG | SF**R**L | ***GVGA*** |
|  | H1.0-B | DSQIKLSIK**R**LV | KS**R**SG | SF**R**L | ***GVGA*** |
| *Homo sapiens* | H1.1 | NNS**R**IKLGIKSLVS | KE**R**GG | SFKL | GTGA |
|  | H1.2 | NNS**R**IKLGLKSLVS | KE**R**SG | SFKL | GTGA |
|  | H1.3 | NNS**R**IKLGLKSLVS | KE**R**SG | SFKL | GTGA |
|  | H1.4 | NNS**R**IKLGLKSLVS | KE**R**SG | SFKL | GTGA |
|  | H1.5 | NNS**R**IKLGLKSLVS | KE**R**NG | SFKL | GTGA |
|  | H1.6/H1t | NNS**R**IKLSLKSLV | QE**R**VG | SFKL | GTGA |
|  | H1.7 | KSG**R**HEAP**R**GQA | THKGL | YF**R**V |  |
|  | H1.8/H1oo | **R**FKYLLKQALATGM**RR** | EQ**RR**G | SFKL | ARGA |
|  | H1.9 | AYHFK**R**VLKGLV | TCKYV | SFTL | GTCK |
|  | H1.10 | G**R**TYLKYSIKALVQN | GE**R**NG | SFKL | GTGA |
|  | H1.0 | DSQIKLSIK**R**LV | KN**R**AG | SF**R**L | ***GVGA*** |
| *Mus musculus* | H1t | NS**R**IKLALK**R**LVN | QE**R**AG | SFKL | GTGA |
|  | H1.0 | DSQIKLSIK**R**LV | KN**R**AG | SF**R**L | ***GVGA*** |
| *Gallus gallus* | H5 | DLQIKLSI**RR**LL | KS**R**GG | SF**R**L | ***GVGA*** |
| *Drosophila melanogaster* | H1 | LAPFIKKYLKSAVV | KE**R**GG | SFKL | GKGA |
